# Experimental and Computational Methods for Allelic Imbalance Analysis from Single-Nucleus RNA-seq Data

**DOI:** 10.1101/2024.08.13.607784

**Authors:** Sean K. Simmons, Xian Adiconis, Nathan Haywood, Jacob Parker, Zechuan Lin, Zhixiang Liao, Idil Tuncali, Aziz M. Al’Khafaji, Asa Shin, Karthik Jagadeesh, Kirk Gosik, Michael Gatzen, Jonathan T. Smith, Daniel N. El Kodsi, Yuliya Kuras, Clare Baecher-Allan, Geidy E. Serrano, Thomas G. Beach, Kiran Garimella, Orit Rozenblatt-Rosen, Aviv Regev, Xianjun Dong, Clemens R. Scherzer, Joshua Z. Levin

## Abstract

Single-cell RNA-seq (scRNA-seq) is emerging as a powerful tool for understanding gene function across diverse cells. Recently, this has included the use of allele-specific expression (ASE) analysis to better understand how variation in the human genome affects RNA expression at the single-cell level. We reasoned that because intronic reads are more prevalent in single-nucleus RNA-Seq (snRNA-Seq), and introns are under lower purifying selection and thus enriched for genetic variants, that snRNA-seq should facilitate single-cell analysis of ASE. Here we demonstrate how experimental and computational choices can improve the results of allelic imbalance analysis. We explore how experimental choices, such as RNA source, read length, sequencing depth, genotyping, etc., impact the power of ASE-based methods. We developed a new suite of computational tools to process and analyze scRNA-seq and snRNA-seq for ASE. As hypothesized, we extracted more ASE information from reads in intronic regions than those in exonic regions and show how read length can be set to increase power. Additionally, hybrid selection improved our power to detect allelic imbalance in genes of interest. We also explored methods to recover allele-specific isoform expression levels from both long- and short-read snRNA-seq. To further investigate ASE in the context of human disease, we applied our methods to a Parkinson’s disease cohort of 94 individuals and show that ASE analysis had more power than eQTL analysis to identify significant SNP/gene pairs in our direct comparison of the two methods. Overall, we provide an end-to-end experimental and computational approach for future studies.

## Background

With the advent of single-cell genomics and the rise of large-scale single-cell RNA-seq (scRNA-seq) studies of hundreds of individuals[1, 2], there has been growing interest in studying how genetic variation impacts gene expression at the single-cell level[2–4]. Such studies could provide new insight into the results of Genome-Wide Association Studies (GWAS), improving fine mapping and helping map key steps from genotype to physiological phenotype, including the identification of cells and cell-specific pathways through which disease genes act. While most studies have focused on expression Quantitative Trait Loci (eQTL) approaches[5], another promising avenue is based on mapping allele-specific expression (ASE) and allelic imbalance. Allelic imbalance analysis is performed by comparing the expression of the maternal and paternal alleles of a given gene within one individual, giving insight into which genes are differentially regulated by Single Nucleotide Polymorphisms (SNPs) on the two alleles. This technique was pioneered for bulk RNA-Seq data and provides an alternative source of information to eQTL analysis[6–8]. Moreover, comparing expression within an individual instead of between individuals should avoid many sources of variability and confounding, potentially increasing power[9, 10].

Compared to bulk ASE, single-cell ASE provides the advantage of understanding heterogeneity across cell types. This approach has been applied successfully to study transcriptional kinetics[11] and dynamic genetic effects on transcription[12]. Statistical methods specifically designed for scASE, such as scDALI[13] and airpart[14], have been developed[15], but have limitations including the inability to account for sample-to-sample variability. A more recent method, DAESC[16], takes into account sample-to-sample variability, though it is specific to the case of allelic imbalance that differs between conditions or along a gradient, rather than testing for allelic imbalance that is consistent in all cells of one type. Moreover, most of these studies have relied on plate-based methods for full-length scRNA-seq, such as Smart-seq2, rather than the more widely used, and far more scalable, 3’ or 5’ end directed droplet-based profiling, though there are exceptions[17, 18]. One main reason has been the assumption that sufficiently long reads and coverage of the entire gene body (from full length scRNA-seq) are needed to recover genetic variants.

There are several advantages of droplet-based snRNA-seq for ASE studies. First, droplet-based methods have the throughput needed for large-scale studies[19]. Second, snRNA-seq has a higher fraction of intronic reads than scRNA-seq[20], which is a potential advantage because there is more genetic variation in introns, thus increasing the ability to identify the allele for a given RNA-seq read. Third, snRNA-seq is compatible with frozen tissues[20], which is crucial in large scale tissue studies from humans – both for collection and for pooling[21, 22]. Finally, snRNA-seq enables a better recovery of diverse cell types [23, 24].

Here, we systematically investigated experimental and computational design choices for their effects on allelic imbalance analysis in droplet-based snRNA-seq data from human samples. We show that many factors, including read length, sequencing technology, and genotyping strategy, can greatly affect the power to detect allelic imbalance. We also developed and explored the use of hybrid selection to target informative regions within genes of interest – a method that greatly improves the power to detect ASE for a given amount of sequencing. We built and tested a suite of computational tools that should enable others to perform ASE analysis on their own sn/scRNA-seq data. Finally, we applied these methods to characterize ASE in human Parkinson’s disease brain samples.

## Results

### ASE analysis pipeline

To measure allelic imbalance, we first built a Nextflow pipeline (Methods) to extract ASE information from scRNA-seq data (Fig. 1a). Although such pipelines now exist for scRNA-seq data, most are built for Smart-seq2 data[25, 26], or do not take advantage of phasing information, instead focusing on each SNP separately[17, 18, 27] (see below for more details). Our pipeline takes in two main inputs for each individual: (**1**) FASTQ files from a transcriptomic assay (compatible with 3’ and 5’ sc/snRNA-seq data, as well as spatial data, and can be easily modified to work with other types of single-cell data) and (**2**) A phased Variant Call Format (VCF) file containing genotype information. These inputs are supplied to STARSolo[28] run with the WASP method[29, 30], which helps avoid biases in ASE, particularly reference bias as discussed below. The output is gene-level expression information and a BAM file annotated with both information from the assay (cell barcode (CBC), Unique Molecular Identifier (UMI), etc.), as well as information about which SNPs each read overlaps. This BAM file is then processed to generate ASE information, both at the SNP level and at the gene level (discussed further below).

**Fig. 1.**
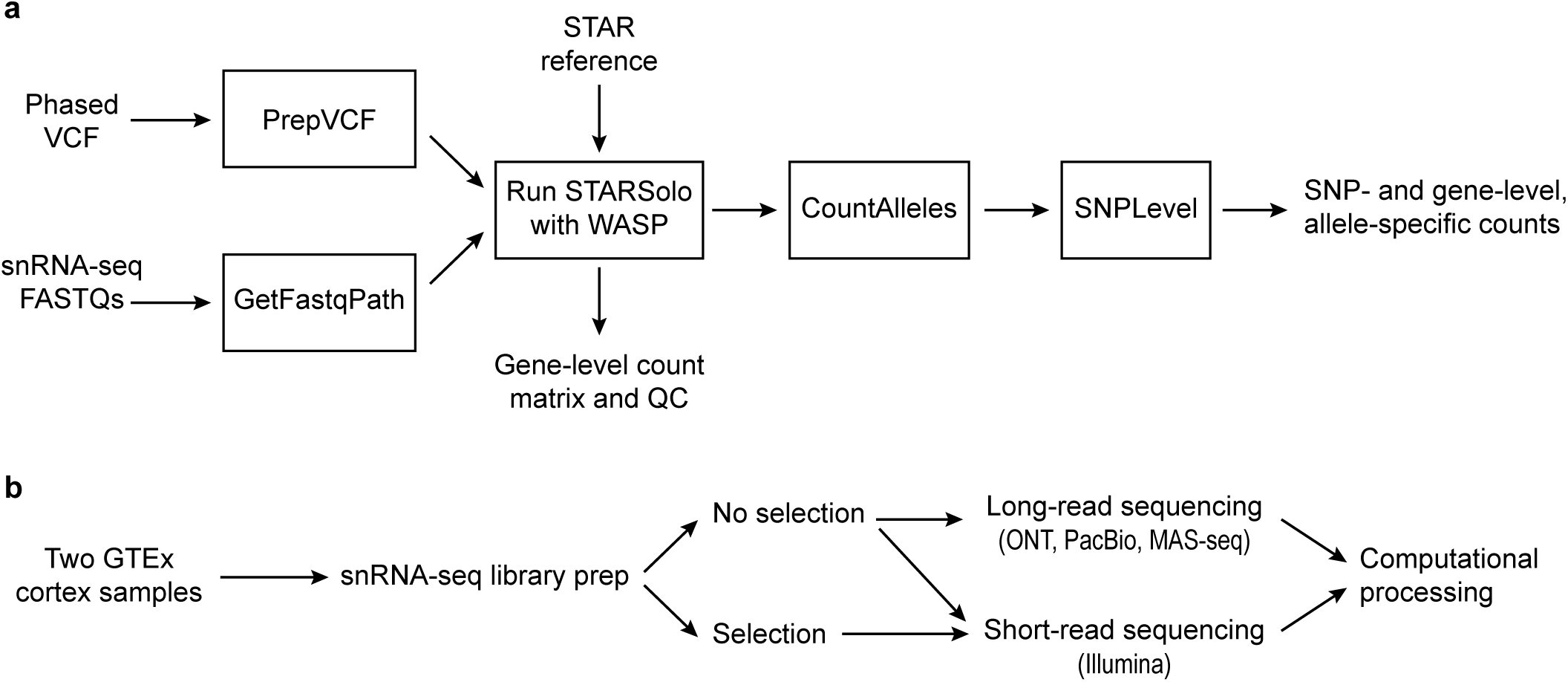
Study overview. **a** Outline of the computational processing pipeline starting from short-read data to generate allele specific expression information (implemented in Nextflow). **b** Sample information and processing for data in Figures 2-4.

### Testing with short-read data

We first analyzed cortical samples from two donors from the Genotype-Tissue Expression (GTEx) project (Fig. 1b). We constructed snRNA-seq libraries from each sample followed by Illumina sequencing (Methods). We accessed a phased genotype VCF file for each individual, previously generated by the GTEx consortium[31]. We processed the snRNA-seq data through our pipeline, resulting in gene expression profiles, ASE information, and sample-level quality control metrics (Supplementary Table 1). The single-nucleus expression profiles were clustered and annotated with cell type labels (Fig. 2a).

**Fig. 2.**
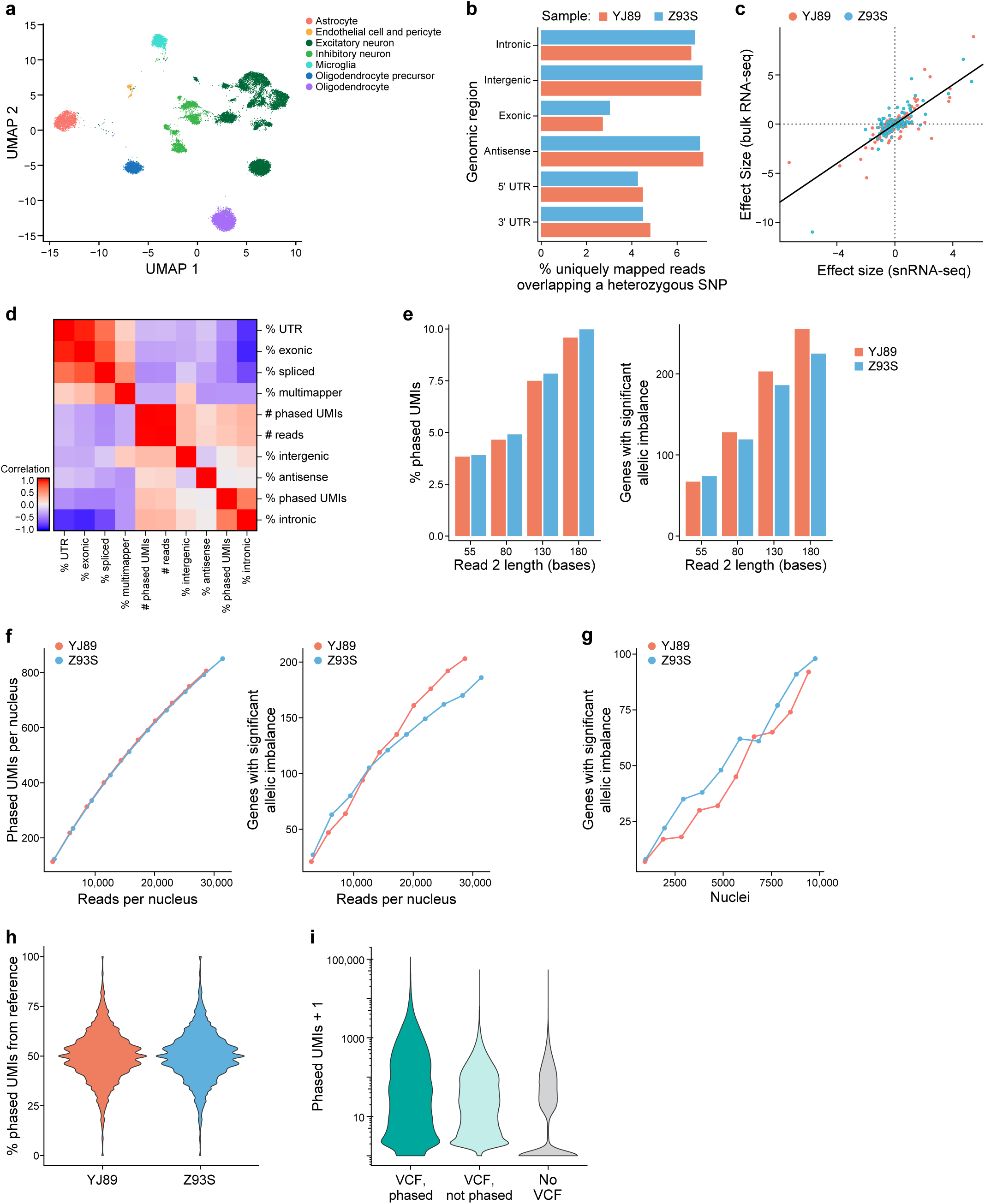
Short-read ASE. **a** UMAP plots of 2 human cortex samples, using 130 base read 2 (see Fig. 2e). **b** Comparison of percentage of uniquely mapped reads that overlap a heterozygous SNP among genomic regions. **c** For each sample and each gene with significant allelic imbalance in that sample, comparison of estimated effect size from our snRNA-seq data to estimate from GTEx bulk data. **d** Heatmap showing the Spearman correlation between droplet-level QC metrics. **e** Effect of trimming read 2 to different lengths on the proportion of phased UMIs (left) and the number of genes with significant allelic imbalance (right). **f** Effect of sequencing depth on the number of phased UMIs (left) and genes with significant allelic imbalance (right); analysis of all nuclei irrespective of cell type. **g** Effect of nuclei number on genes with significant allelic imbalance. **h** Violin plots showing for each SNP the percentage of UMIs that map to the reference allele. **i** Violin plots for each gene with at least one phased UMI showing the maximum number of phased UMIs over all SNPs in that gene, with phased VCF, unphased VCF, or no VCF (genotypes estimated from the snRNA-seq data). Without phasing information, it was not possible to combine reads overlapping different SNPs in the same gene into one gene-level measure because the genotype of each SNP is on each allele was unknown. Instead, the maximum number of phased UMIs over all SNPs in that gene is reported, which can be considered as a of measure the maximum power to detect allelic imbalance in that gene.

The genomic region of the read (e.g., intronic *vs*. exonic) affected our ability to extract ASE information. It is only possible to extract ASE information from reads that overlap a heterozygous SNP because such overlap is required for assigning the allele of origin to a read. As such, for each sample we looked at the percentage of reads from each region that overlapped a heterozygous SNP (Fig. 2b). The vast majority of reads did not overlap such SNPs, so that they were not usable for ASE. Their proportion, however, differed from region to region, with exonic reads having the smallest fraction overlapping heterozygous SNPs, while intronic, antisense, and intergenic SNPs all had a higher fraction, supporting our hypothesis. Given the higher level of intronic reads in single-nucleus *vs*. single-cell RNA-seq data[20], there may be more power for allelic imbalance in snRNA-seq data, at least on a per-read basis, though a more direct comparison would be needed to confirm this.

As another control, we compared our gene-level ASE results to previously published bulk RNA-seq for these two samples – obtaining the data from the GTEx portal[31]. As expected, there is high agreement (Spearman correlation = 0.68, Pearson correlation = 0.80; Fig. 2c) – providing confidence in the output of our single-nucleus ASE pipeline.

Next, we calculated droplet-level QC metrics for each cell barcode in a library, including percentage intronic reads, percentage antisense reads, etc. For ASE analysis, we focused on “phased UMIs,” which are those UMIs that we can assign to an allele and use to calculate ASE. We calculated the correlation of the droplet-level metrics with the number and fraction of phased UMIs for each cell (Fig. 2d). As expected, the total number of reads is highly correlated with the total number of phased UMIs, and the percentage intronic reads is highly correlated to the percentage of UMIs that are phased. Similarly, there is a higher proportion of phased UMIs when we include intronic reads in our analysis (Fig. S1a). For each gene with significant ASE, comparing the allelic imbalance effect size between the two approaches with introns and without introns, however, there was very high agreement overall (Fig. S1b).

We next explored the role of read length in generating ASE information. Intuitively, the longer a read is, the more likely it is to overlap a heterozygous SNP, so that a larger fraction of longer reads than shorter reads should be usable for ASE analysis. To assess this, we trimmed read 2 (the cDNA read) from the 3’ end to different lengths, ranging from 55 bases to 180 bases, processed the resulting data with our pipeline, and calculated the percentage of phased UMIs. The higher this percentage, the more power we will have per UMI. As expected, longer reads yielded a higher proportion of phased UMIs (Fig. 2e) and allowed us to detect more allelically imbalanced genes in aggregated data from all cells (Fig. 2e) or from most cell types individually (Fig. S2). This must be balanced with the higher cost per read for longer Illumina reads (so one is likely to be able to afford more reads with shorter reads, something that also affects the power, see below), with the choice depending on many experiment-specific details. Here, we focused on the case with 130 base read 2, which corresponds to 150 cycles of sequencing.

Building on the above, we considered how the number of reads sequenced affected the number of phased UMIs. As expected, deeper sequencing coverage leads to the identification of more phased UMIs and more allelic-imbalanced genes (Fig. 2f). Though the rate of increase slowed down as the number of reads sequenced increased, it did not plateau for either metric. Similarly, as the number of cells increased, the power to detect allelically imbalanced genes increased (Fig. 2g), and it did not plateau with ∼10,000 nuclei per sample.

One common issue with ASE is reference bias[29, 32]: since reads containing the reference allele can align at a higher rate to the reference genome than those with the alternative allele, this can lead to biases in read mapping and false positive allelic imbalance results. Although our pipeline mitigates this problem, by using the WASP [29] implementation in STAR[30], we explored the effects of reference bias in our data. We looked at the number of phased UMIs for each SNP, calculated the percentage mapping to the alternative allele, and plotted their distribution for all SNPs with high enough expression (Fig. 2h). For both individuals, these distributions were centered at 50%, consistent with at most small levels of reference bias.

### Alternative approaches without phasing information

While the above analysis uses phased genotype data to share information between SNPs on the same haplotype (covered by different reads), an alternative approach is to use only the reads directly overlapping a given SNP. For bulk RNA-seq data, the use of phasing information has been shown to improve power[6] and to enable allelic imbalance to be derived for SNPs that might not be covered by any reads, such as intergenic SNPs. However, using phased genotypes can lead to incorrect ASE estimates if there are errors in the phasing, and can be problematic for genes with differential ASE in different RNA isoforms. Fortunately, current population-based phasing approaches work well at the genomic distances (gene length) required for *cis*-eQTL analysis[33, 34]. In the case of scRNA-seq data, however, we often lack a phased VCF and might not even have genotype information at all. As such, some scRNA-seq ASE pipelines do not use phasing information, and (by default) use the scRNA-seq to estimate a genotype for each sample[18]. Such an approach expands ASE analysis to many more datasets, but limits the SNPs that can be explored. Moreover, a large allelic imbalance at a given SNP could, at least in theory, lead to issues with genotyping because the less expressed allele might be hard to identify.

To explore approaches that do not use phasing information, we generated ASE analyses with three different methods: (**1**) using the phased VCF, (**2**) only counting reads directly overlapping a given SNP in our VCF (no phasing information), and (**3**) calling SNPs directly from the snRNA-seq data and only counting reads directly overlapping a SNP identified by the snRNA-seq (no genotype information). In theory, it might be possible to phase genotype data derived from snRNA-seq[35], but to our knowledge this has not been demonstrated for droplet-based snRNA-seq. For the last approach we only considered known SNPs in the population – thus excluding *de novo* variants, given the risk of sequencing errors[36, 37]. Using the phased information yielded the most phased UMIs per gene, with the worst performance being the analysis with no VCF (Fig. 2i) – noting that the exact performance of this last method will depend on the method used to call genotypes from snRNA-seq.

### Transcript-level analysis

Our analysis methods to this point have explored the effects of a given SNP on the expression of a given gene, but a given SNP might differentially affect each of the transcripts of a given gene and our standard approach cannot distinguish such effects. Moreover, recent studies have shown that taking into account isoform-level information can improve power for eQTL analysis in bulk RNA-seq data[38]. Although the 3’-biased, short-read nature of our droplet-based snRNA-seq can make it difficult to extract isoform-level information, there are published methods that attempt to explore isoform-level changes with 3’-biased scRNA-seq, particularly changes in the 3’ UTR including alternative polyadenylation sites[39, 40]. We explored one such method known as Sierra, which identifies peaks of expression within genes, i.e., regions of the gene with a high density of reads[41]. If cDNA priming occurred only at the poly(A) tail of mRNAs, these peaks would correspond to different 3’ ends of transcripts, but due to “internal priming” from other parts of the transcript, the peaks cover more than just the 3’ end of the transcripts. Peaks from internal priming, however, provide information about splicing because a peak in a given exon or intron indicates that exon or intron is transcribed from the gene. As such, we modified our pipeline to report ASE information at the level of the peaks returned by Sierra (Methods).

Of the peaks identified by Sierra (Fig. S4a), 79.5% were in introns and 28.9% were in exons – with some overlap between the different classes, thus adding up to >100% (Methods). Only 5.9% of peaks crossed a splice junction.

To test our peak-level ASE pipeline, we analyzed snRNA-seq data from both GTEx samples with Sierra, producing sample-specific sets of peaks. We then used Sierra to combine these peaks between the samples, resulting in a merged set of peaks, which include putative 3’ UTR regions, for downstream analysis and for use with our pipeline. This resulted in ASE UMI counts for each peak in each cell. We next performed ASE analysis at the peak level – using the same analysis as for the gene level, except with the peak-level ASE information. Compared to the gene-level approach, the peak-level analysis resulted in slightly fewer genes with ASE, with a fair amount of overlap between the genes identified at the peak level and at the gene level (Fig. S4b), such that 32.3% of genes with significant imbalance had at least one significant peak and 38.6% of peaks with significant imbalance were in a gene with significant imbalance (Supplementary Table 2).

### Long-read sequencing

Because longer Illumina reads produced more phased UMIs (Fig. 2e), we explored long-read sequencing technologies as an alternative approach. In particular, we started from the unfragmented RT-PCR cDNA products from the 10x Chromium libraries for the two GTEx samples (see above), generated libraries appropriate for each of the long-read technologies, and sequenced them with the Sequel II (Pacific Biosciences, PacBio), GridION (Oxford Nanopore Technologies (ONT)), or PromethION (ONT) platforms (Supplementary Table 3). In addition, we tested MAS-ISO-seq (MAS-Seq), which combines cDNA concatenation with PacBio sequencing (either with the Sequel IIe or Revio), resulting in sequencing many more cDNAs than with a standard PacBio run[42], using both the Sequel II and the newer Revio instruments. To overcome higher sequencing error rates for ONT-based sequencing, we applied the R2C2 method[43] to construct libraries as it enables multiple passes of sequencing of the same original DNA. PacBio and MAS-Seq sequence the same DNA with multiple passes, so that an approach such as R2C2 is not necessary. These data were processed with existing pipelines[36, 42] (Methods) to collapse together multiple sequencing passes to create a consensus sequence, split concatenated reads into their corresponding cDNAs in the case of MAS-Seq, align these reads to the genome, and annotate the resulting BAM files with 10x Chromium-specific information, most notably CBC and UMI. We then annotated each read with a list of any genes that read overlapped and extracted ASE information from these BAM files (Methods). The subsequent analysis described here used only these long reads processed as described above.

We first investigated the length of these reads. In particular, since each read includes information about the UMI, CBC, poly(A) region, and adapter sequences, we looked at the length of the region mapping to the genome. Although there are slight differences among the technologies, most had a median length of 800-1000 bases comparable to a previous study[44], though MAS-Seq was slightly shorter (Fig. 3a). This is much longer than with Illumina sequencing, though still not long enough to span the entire length of most mRNA and pre-mRNA molecules.

**Fig. 3.**
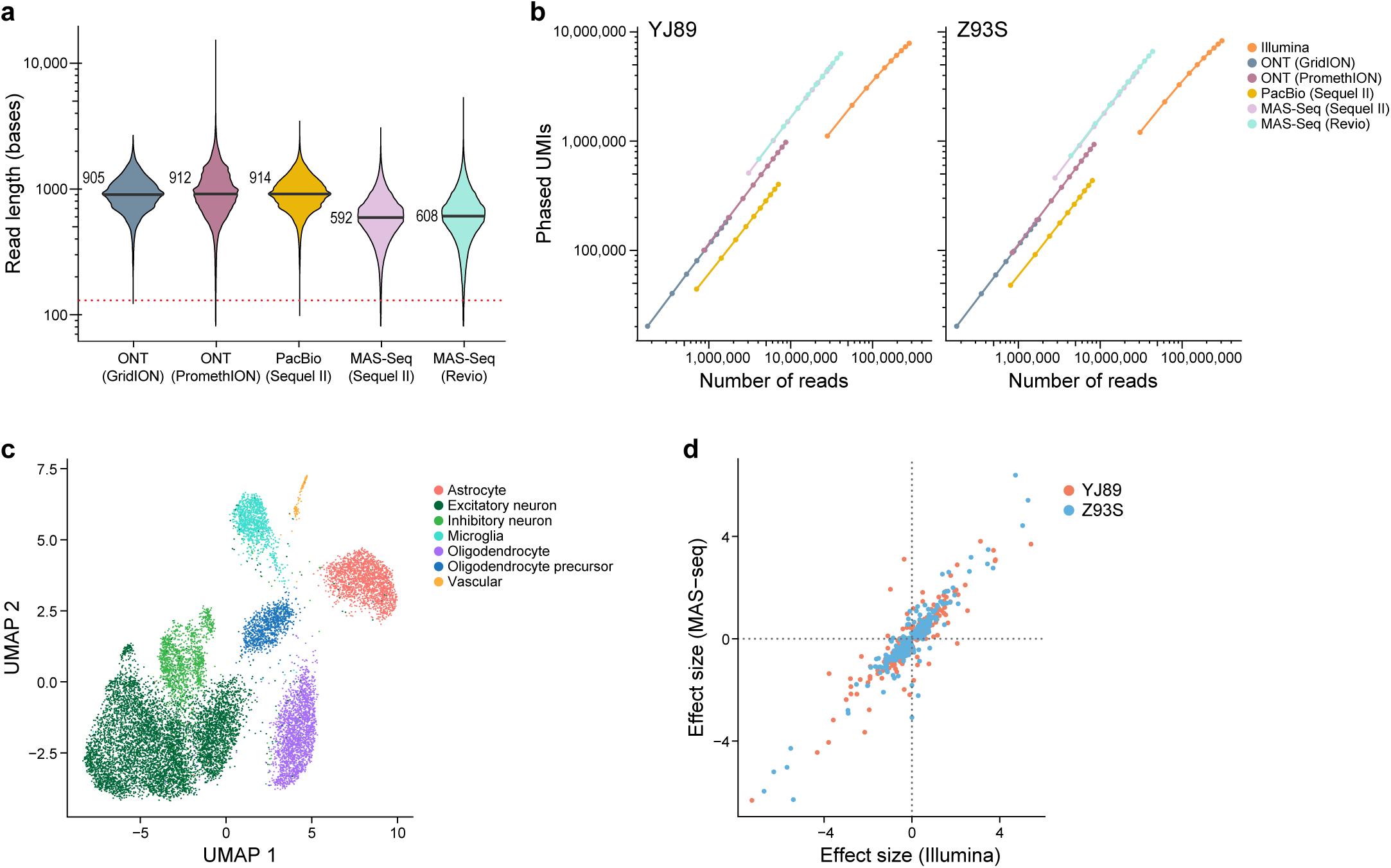
Long-read ASE. **a** Violin plots of the length of reads uniquely mapped to the genome, with median shown. Dotted line shows Illumina 130 base reads. **b** Downsampling analysis of phased UMIs *vs.* sequencing depth. **c** UMAP of the MAS-Seq data. **d** Comparison of the estimated allelic imbalance effect size in Illumina and MAS-seq for genes that are significantly allelically imbalanced in at least one of the two datasets.

One of the main concerns historically with long-read technologies for genotyping-based applications has been their high error rates. To investigate this, we looked at the frequency of mismatches, insertions, and deletions in each technology relative to the reference genome, scaled by the number of aligned bases (Fig. S3). We note that some of these mismatches were due to polymorphisms in the genome, so the numbers reported here are an upper bound on the actual error rate. Illumina had very few insertions and deletions (0.00015 - 0.00021 per matching base), while ONT-based technologies had many more (0.011 - 0.016 per matching base), and PacBio and MAS-Seq were had intermediate numbers (0.0009 - 0.0036 per matching base). For single base mismatches, Illumina and ONT had similar frequencies (0.011 - 0.012 per matching base pair for Illumina, 0.007 - 0.010 per matching base for ONT), while PacBio and MAS-Seq had substantially fewer (<0.0026 per matching base). Consistent with previous reports[37, 42, 45], PacBio and MAS-seq had a fairly low error rate, with more indels but fewer mismatches than Illumina, and ONT-based methods had a higher error rate.

As expected, for a given number of reads, the long-read technologies recovered many more phased UMIs than Illumina with a 130 base read 2 (Fig. 3b). Due to the greater number of reads recovered with Illumina, however, we still obtained more phased UMIs overall with Illumina than with the long-read technologies except for MAS-seq with Revio, which had a comparable number. This indicates that MAS-seq and Illumina are both powerful methods for performing gene-level ASE analysis of snRNA-seq data. Our estimates suggest for every MAS-seq segmented read (S-read) one would need to sequence ∼5.5 Illumina reads (with 130 bases in read 2) to get the same number of phased UMIs though the exact number depends on the level of saturation, so that MAS-seq would be more cost effective than Illumina if the cost of one S-read in MAS-seq is less than seven times the cost of an Illumina read.

We also compared the per gene allelic imbalance effect estimates in Illumina and MAS-seq (Revio), which was the most efficient long-read technology (Fig. 3b). We identified the expected cell types with MAS-seq (Fig. 3c). In an analysis limited to genes that are significant in at least one of the two technologies, we see high agreement in terms of effect size (Pearson correlation 0.91, p-value < 2.2e-16; Fig. 3d).

### Isoform analysis of long-read sequencing

One advantage of long-read sequencing is the improved ability to assign reads to particular isoforms. As such, we constructed a pipeline to extract isoform-level ASE information from long-read data mapped to the genome (Methods). We assigned each read either to a spliced isoform from an existing reference or to an unspliced isoform, which would likely be derived from pre-mRNA, based on the overlap between each read and each isoform (Methods). Reads that could not be assigned to a specific isoform in this way were discarded from downstream analysis and could derive from either mRNA or pre-mRNA. Using the MAS-Seq data (Revio), we were able to assign 65.2% of gene-level phased UMIs to either a spliced or an unspliced isoform. This, however, was largely driven by unspliced isoforms, with only 4.3% of the isoform-level UMIs assigned to spliced isoforms. For phased UMIs, only 3% were assigned to spliced isoforms. Consistent with our intron *vs*. exon analysis in short reads (Fig. 2b, Fig. S1a), a higher percentage of UMIs were phased in unspliced isoforms than spliced isoforms (25.0% *vs.* 17.4%). The vast majority of phased UMIs were assigned to isoforms from pre-mRNA for most genes (Fig. S4c). On a per-nucleus basis (Fig. S4d), there were relatively few (11 on average) phased UMIs assigned to spliced isoforms, with many more phased UMIs (450 on average) assigned to unspliced isoforms. Still, there are many spliced isoforms with moderate expression – 2,124 spliced isoforms with >10 phased UMIs in at least one of the two samples and 492 spliced isoforms with >10 phased UMIs in both samples. It may be possible to improve the alignment of reads to transcripts with an extended transcriptome reference, which is common in long-read RNA-seq analysis and particularly important in 3’ scRNA-seq[46, 47].

We extended this analysis using the above genome-based pipeline with slight modifications (Methods) to the GTEx snRNA-seq short-read data. With varying lengths for read 2, there were very few phased UMIs for spliced transcripts (1-2 per cell), but hundreds for the unspliced transcripts (Fig. S4e). This was much fewer spliced transcripts per cell than for the MAS-Seq data (Fig. S4d) – suggesting MAS-Seq has a major advantage over short-read data for isoform-level ASE analysis.

### Targeted sequencing

In many research studies, it may be optimal to generate ASE information for a specific subset of genes rather than the entire transcriptome. For example, in a study focused on GWAS loci for a particular disease, it could be best to target genes near those GWAS loci. Similarly, we would prefer to focus our sequencing near heterozygous SNPs, since those are the only reads that can be used to estimate ASE. Enriching for reads in those regions should provide more power to detect ASE in those genes, without expending resources sequencing genes that are not relevant to the study, as well as reducing both the computational resources required and the level of multiple hypothesis correction.

To test this, we used hybrid selection[48] to select within our two GTEx snRNA-seq Illumina libraries for cDNAs containing regions of interest. We selected a list of 100 genes of interest to have a variety of properties—from long to short, highly expressed to lowly expressed, cell type-specific and more widely expressed, and known eQTLs *vs.* not (Methods). We further identified regions to target by extracting regions in these genes that had coverage in our snRNA-seq Illumina data and were near a heterozygous SNP in one of our two samples. We then designed baits to target these regions, performed a hybrid selection experiment with the baits and the snRNA-seq libraries, sequenced with Illumina (202 bases for read 2) to just under 50,000 reads per cell as reported by Cell Ranger, and processed the data with our ASE pipeline (Methods). We also resequenced the non-selected library in the same sequencing run to avoid confounding due to batch variability, read length, and sequencer used.

To assess the extent of the enrichment, we analyzed the phased UMIs in the targeted genes. Data from the hybrid-selected libraries have a much higher proportion of phased UMIs aligned to the selected genes, though that proportion decreased as sequence coverage increased (Fig. 4a). To explain this decrease, we looked at each UMI in the hybrid-selected data and asked how many reads supported that UMI. For UMIs in targeted genes, there was a median of 22 reads per UMI and a mean of 84.6 reads per UMI, while the non-targeted genes had a median of 1 read per UMI and a mean of 1.65 reads per UMI (Fig. 4b), suggesting that we sequenced more deeply than needed for the targeted genes in the hybrid-selected data.

**Fig. 4.**
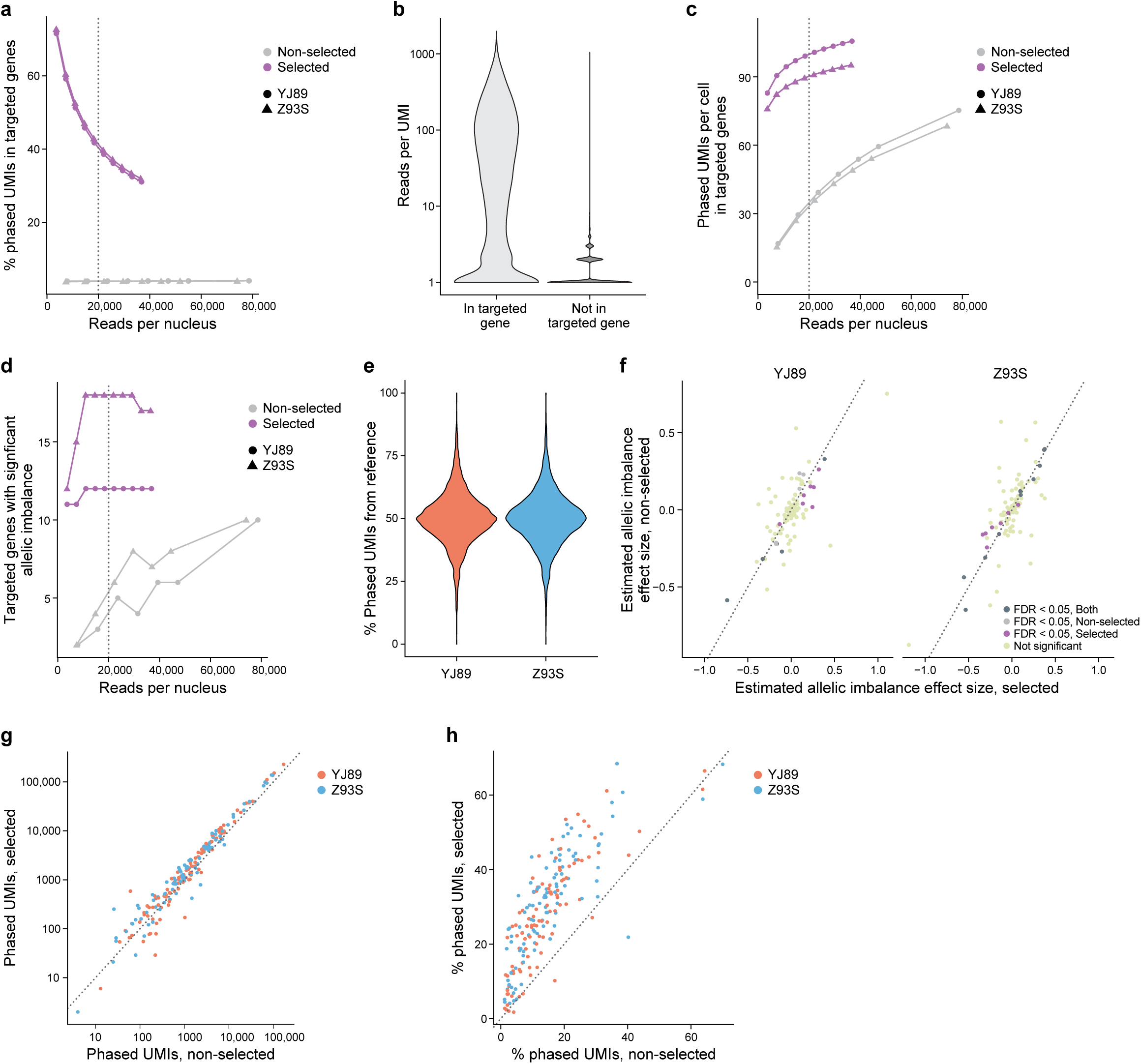
Hybrid selection. **a** Proportion of phased UMIs in one of the targeted genes with or without selection at different sequencing depths. **b** Violin plots of the number of reads in selection data supporting each UMI in targeted and non-targeted genes. **c** Phased UMIs recovered in the 100 targeted genes at varying sequence depths with and without selection. **d** Number of genes with significant ASE among the 100 targeted genes at varying sequence depths with and without selection. Dotted line in **a**, **c,** and **d** indicates 20,000 reads per call, vendor recommended sequencing depth. **e** Violin plots of the percentage of UMIs overlapping each SNP mapping to the allele in the reference genome in the selected data. **f** Comparison of the estimated allelic imbalance of the selected genes in the selected and non-selected data using all reads. **g** Comparison of the number of phased UMIs in targeted genes in the non-selected data *vs.* selected data. **h** Comparison of the percentage of UMIs phased in each targeted gene in selected *vs.* non-selected data. Dotted line in **g** and **h** represents the line x = y.

In addition to looking at the proportion of phased UMIs from targeted genes, we examined the total number of phased UMIs from the targeted genes (Fig. 4c). Even at fairly low levels of sequencing (<3,700 reads per cell), the hybrid-selected data had many (>76) phased UMIs per cell in the targeted genes, though this started to plateau with additional sequencing coverage. By contrast, for the non-hybrid-selected data, much more sequencing was required to reach the same level of phased UMIs in the targeted genes (<76 phased UMIs per cell even with >74,000 reads per cell, more than 20x as many reads). Similarly, even at the lowest sequencing depth tested we were able to detect 11-12 allelically imbalanced genes among the 100 targeted genes in the hybrid-selected data, but much more sequencing was required for the same power in the non-hybrid-selected data which never had more than 10 significant genes even with 20 times more coverage (Fig. 4d). As might be expected, many more of the genes with known eQTLs are found to be allelically imbalanced than those without known eQTLs in both hybrid-selected and non-hybrid-selected data (Fig. S5a). Analysis at a finer level revealed that with selection reads were much closer to the baits (Fig. S5b), with ∼80% of uniquely mapping reads overlapping a bait in the selected data, but >90% of uniquely mapping reads being >10,000 bp away from the closest bait without selection.

We checked whether our hybrid selection approach biases the data to the reference allele, leading to false positives in a similar way to reference bias. Although at least one previous study[49] has suggested such biases are minimal in the case of DNA sequencing, we decided to check this in our own data, by several lines of evidence. First, there was a distribution centered around 50% of phased UMIs deriving from the reference allele for each sample (Fig. 4e), indicating that there is no evidence of reference bias in the per SNP ASE estimates in our data. Second, the estimates of allelic imbalance for each gene between the hybrid-selected and non-hybrid-selected data (using all reads in both cases) are highly concordant between the two assays (Pearson correlation of 0.75, p-value < 2.2*10^-16^; Fig. 4f), consistent with minimal bias in the selected data. Third, only six of 18,770 SNPs in the 100 targeted genes (0.03%) had significantly different (FWER < 0.05, Fisher’s exact test) ASE levels between selected and non-selected data for either sample, whereas for SNPs not in the selected genes it was six of 182,854 (0.003%). Thus, a SNP in one of the 100 targeted genes was about 10 times as likely to show significant difference between the selected and non-selected data than other SNPs, though this was confounded by the SNPs in the targeted genes having much higher coverage in the selected data than those in genes that were not targeted. Importantly, there does not seem to be a bias towards the reference allele, with 54.5% of the 100 SNPs with the smallest p-values having increased expression of the alternate allele *vs.* reference in the data with selection relative to the data without selection.

Notably, the six SNPs with significant differences between selected and non-selected had a higher percentage of ambiguous UMIs. An ambiguous UMI is a UMI/CBC pair with different reads that assign the same SNP or gene to different alleles. Our pipeline counts such UMIs, but instead of assigning them to an allele they are assigned as “ambiguous” and not used in downstream analysis. This modest increase in ambiguous UMIs might result from higher rates of PCR recombination[50] due to additional PCR cycles needed for hybrid selection. Such reads accounted for only 0.32% of phased UMIs in the data without selection, while accounting for 1.58% of phased UMIs in the data with selection. In addition, with selection there were more ambiguous UMIs in the SNPs in targeted than non-targeted genes (3.8% *vs.* 0.13%). Moreover, the level of ambiguous UMIs was much higher (18.1%) for those SNPs in targeted genes with ASE differences between selected and non-selected data —raising the possibility that the removal of the ambiguous reads led to higher variability than expected by the Fisher’s exact test, which might have led to significant results in the above analysis, though this remains uncertain. In light of these findings, it might be worthwhile to implement alternative filters[50] to help reduce the effect of ambiguous UMIs in addition to or instead of the filter in our pipeline.

We further looked at how well the method performs at the gene level. As expected[48], the number of phased UMIs for each gene with and without hybrid selection were very highly correlated (Fig. 4g, Spearman correlation = 0.97), suggesting that expression before selection is a strong predictor of the expression after, with more uncertainty for the lowly expressed genes. Furthermore, most genes had more phased UMIs with hybrid selection than without (Fig 4g). For each gene, we also compared the enrichment (total number of phased UMIs with selection divided by the total number of phased UMIs without selection) to QC metrics from the data without selection (Fig. S5c). There is no evidence of a strong correlation between enrichment and any of these QC metrics. Genes with lower expression, as measured by percentage expressing cells, total UMIs, etc., show less consistent enrichment than those expressed at higher levels. Similarly, there was not a strong association between the reason genes were chosen and the enrichment score (Fig. S5d).

Within the targeted genes, we used hybrid selection to target regions near heterozygous SNPs to increase the percentage of UMIs that are phased. In our experiments, the percentage of UMIs that are phased in the hybrid-selected data is greater than that in the non-hybrid-selected data for almost all targeted genes (Fig. 4h).

The above analysis suggests that hybrid selection should increase power for ASE analysis in regions and genes of interest while requiring much less sequencing resources.

### Parkinson’s disease SNP-level analysis

We next sought to extend our findings to a larger number of individuals. Allelic imbalance is often studied in data from many individuals[6, 16] because determining which SNPs are driving allelic imbalance, requires data from many individuals to tease apart the effects of a given SNP since many SNPs are on each haplotype, any one of which could be driving allelic imbalance.

To this end, we used data from a Parkinson’s disease (PD) study consisting of snRNA-seq from the mid-temporal gyrus (cortex) of 94 individuals (Lin et al., manuscript in preparation) with clustering and cell type annotations reported elsewhere (Fig. 5a; Methods). The donors consist of about equal numbers of healthy controls, individuals with PD, and individuals with incidental Lewy body disease (ILBD; pathology without obvious clinical symptoms).

**Fig. 5.**
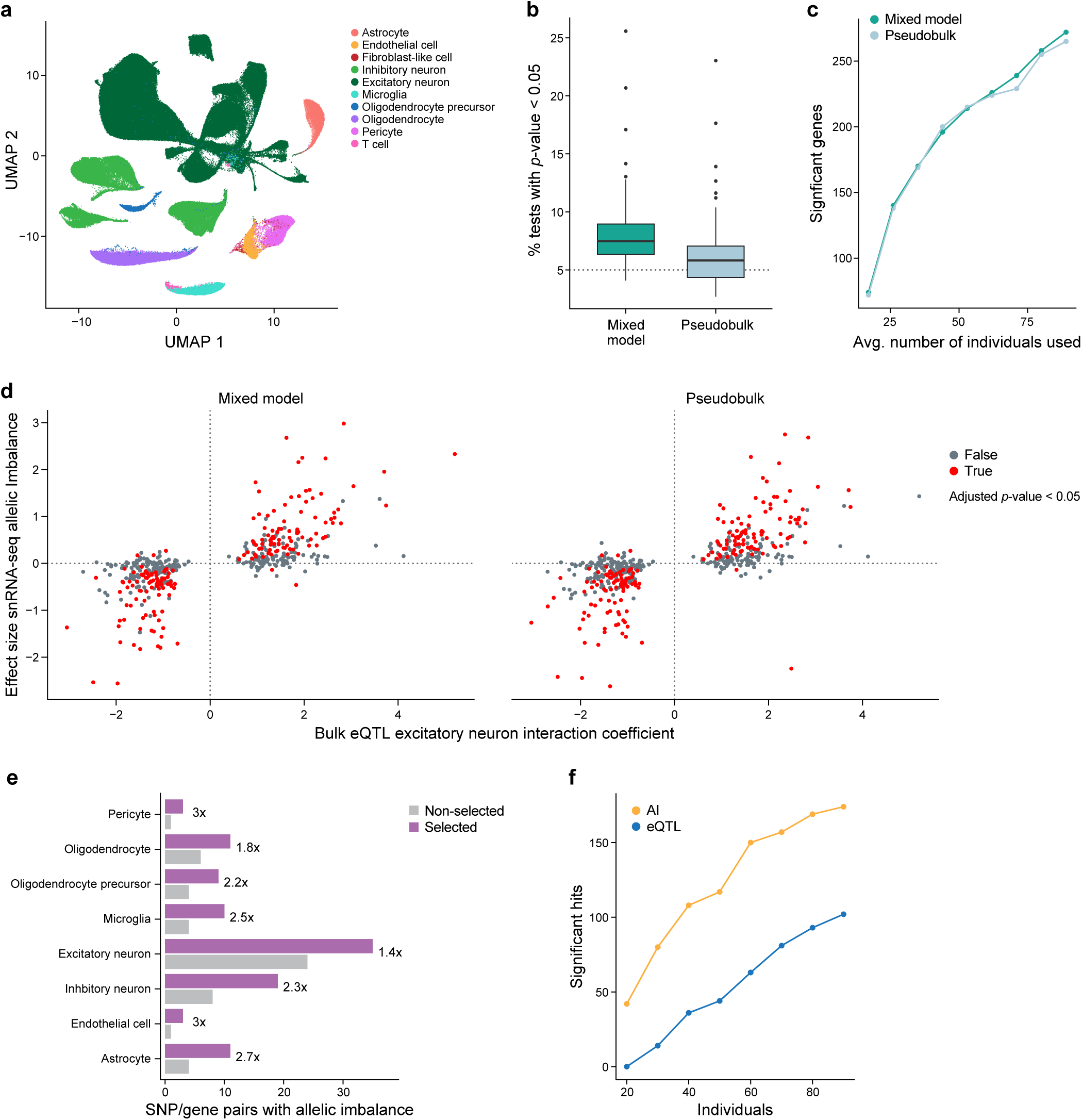
Parkinson’s disease ASE. **a** UMAP of PD mid-temporal gyrus snRNA-seq data. **b** Box plot of false positive rate (FPR) analysis. Results of random permutation (100 times, Glutamatergic neurons data) of the alternate *vs.* reference allele for each individual to generate a dataset with no true allelic imbalance. Shown for each permutation is the percentage of comparisons with p-value < 0.05, which should be around 5% if the method controls the FPR. Boxplots denote the medians and the interquartile ranges (IQRs). The whiskers of each boxplot are the lowest datum still within 1.5 IQR of the lower quartile and the highest datum still within 1.5 IQR of the upper quartile. **c** Evaluation of methods to detect allelic imbalance for known cis-ieQTLs with downsampling of individuals. **d** Comparison of the estimated allelic imbalance effect size in the snRNA-seq data for excitatory neurons with our ASE methods to the excitatory neuron interacting eQTL effect size from published bulk RNA-seq data. **e** Bar plot comparing the number of significant allelic imbalanced SNP/gene pairs for targeted SNPs in each cell type. **f** Comparison of significant SNP/gene pairs detected by eQTL or allelic imbalance analysis with different number of individuals sampled. Significant hits had a Bonferroni-corrected p-value < 0.05.

Previous studies on allelic imbalance in single-cell and single-nucleus RNA-seq data have mostly analyzed individual samples [13, 14]. DAESC, a recent method for studies with multiple samples[51] relies on differential allelic imbalance, comparing ASE levels between conditions. In our study, however, we focus on allelic imbalance in a particular group of cells regardless of condition. To test this we implemented two methods, both based on the beta-binomial distribution (Methods). One is a pseudobulk method, where the reads from cells of the same type from the same individual are aggregated and tested with a standard beta-binomial regression model from the aod package, and the other is a mixed model method using the glmmTMB package[52], where cells are aggregated, but instead there is an extra random effect term in the analysis to account for individual to individual variability, the same underlying model tested in DAESC. Similar methods are used for differential expression between conditions in single-cell data except with different underlying noise models[53, 54].

To assess how well these methods control the false positive rate, we randomly permuted the reference and alternative allele for each individual and gene in excitatory neurons (Methods) to create a dataset that should have no significant allelic imbalance. We then used both pseudobulk and mixed model methods to test for allelic imbalance. We repeated this 100 times and looked at, for each permutation, the proportion of tested SNPs with p-value < 0.05. We would expect this to be ∼5% in a well-calibrated method. In these analyses, the pseudobulk-based method has a median value close to 5%, though with high variability between permutations, indicating it did well in controlling the false positive rate, while the mixed model more frequently has SNPs with p-value < 0.05 indicating it generated a higher than desired false positive rate (Fig. 5b).

In addition, we examined the power of each method. Estimating power can be challenging due to the lack of true positives and, as such, sometimes simulated data are used. Instead, we used actual data and counted how many significant allelic imbalance events were detected among SNP/gene pairs that were likely to be true positives based on previous evidence. In particular, we took a set of SNP/gene pairs that were identified in a previous study[55] as excitatory neuron *cis*- interacting eQTLs (cis-ieQTLs), which are bulk eQTLs whose effect scales with the estimated proportion of excitatory neurons,. These ieQTLs were identified by applying deconvolution-based methods to bulk cortical tissue to calculate the proportion of cells in that sample that are excitatory neurons and using those estimates to find eQTLs where the eQTL effect size differs with the proportion of excitatory neurons. These SNPs have been proposed to have stronger effects on the expression of the gene of interest in samples with more excitatory neurons, so that they are strong candidates to be cis-eQTLs in excitatory neurons, with caveats such as differences in cellular contents (RNA from the whole cells *vs.* nuclei) and RNA-seq library construction methods. We sampled different numbers of individuals from the entire dataset and tested for allelic imbalance with both pseudobulk and mixed model methods. As expected, with increasing numbers of individuals, more significant allelic imbalance events were detected (Fig. 5c). Both methods detected many allelic imbalance events, though the mixed model methods did tend to detect slightly more at a given FDR cutoff.

Notably, there was very high agreement between the expected effect size from the snRNA-seq data, using both the mixed model and pseudobulk approach, and that in the bulk RNA-seq data (Fig. 5d), particularly among the significant results. This shows that allelic imbalance analysis of snRNA-seq can recover known biological signals. Although there are a sizeable number of SNP/gene pairs that are significant in the bulk but not in the single-nucleus data, this is not surprising given the differences between single nuclei and bulk, that not all ieQTLs are necessarily eQTLs in excitatory neurons, and that the bulk results are from thousands of individuals whereas the snRNA-seq data are from fewer than 100.

### Parkinson’s disease gene targeting

To extend our findings with ASE analysis of PD data, we applied hybrid selection, as with the two GTEx samples, to target 101 SNP/gene pairs identified as eQTLs in PD GWAS loci (Methods). We generated sequence data targeting relevant regions of these genes for 91 individuals in our PD dataset. In general, the selected libraries were sequenced less deeply than the non-selected libraries, as measured by reads per cell for each individual, (Fig. S6a), with a median of 42% of the sequencing depth in the selected data compared to the non-selected data and with 86 of 91 samples having more sequencing in the non-selected data than the selected data. Despite this, the selected data have much higher saturation than the non-selected data (saturation of 50.4% *vs.* 15.7% as reported by STARSolo).

For each cell type, we performed our pseudobulk-based allelic imbalance analysis of both the selected and non-selected data and compared the number of SNP/gene pairs with significant allelic imbalance in each cell type (Fig. 5e, Supplementary Table 4). Despite having fewer reads per cell in almost all samples, the selected data have many more significant hits than the non-selected data (1.56x (excitatory neurons) to 3x (endothelial cells)), with 53 SNP/gene pairs that were significant in the selected data but not in the non-selected data, but only four SNP/gene pairs that were significant in the non-selected data but not in the selected data. We note that 19 of the 53 SNP/gene pairs were not tested in the non-selected data due to low sequence coverage of those genes. To investigate in a more systematic way, we determined the number of significant SNP/gene pairs in each cell type after downsampling the selected data to a depth of 5,000 reads per cell (Methods, Fig. S6b,c). There were very similar results to those without downsampling, consistent with the plateau we observed with increasing sequence coverage for gene-level allelic imbalance in the GTEx data with selection (Fig. 4c,d, Supplementary Table 4). Our results suggest the selected data have more power than the non-selected data to detect allelic imbalance at the SNP level, even with much less sequencing coverage.

There is very high agreement in the estimated effect size between the selected (without downsampling) and non-selected data (Fig. S6d) (Pearson correlation = 0.86, p-value < 2.2*10^-16^). This suggests both experiments detected the same signal and we had more power to detect it in the selected data.

### Isoform-level analysis

In addition to using ASE to find SNPs that affect gene expression, we were also interested in the effects of specific SNPs on levels of different RNA isoforms of a gene. To do this we built on the Sierra-based methods[41] discussed above. We used Sierra to call merged peaks from the PD non-selected dataset and processed these results with our pipeline to generate peak-level ASE information.

First, we tested how many peaks had significant ASE in excitatory neurons (Fig. S7a). As expected, as the number of individuals increases, we observe many more significant peaks. We also checked for peaks in which the ASE ratio in that peak differs from the ASE ratio of the rest of the gene (Methods), which is similar to differential transcript usage (DTU), and observed similar results (Fig. S7b *vs*. Fig. S7a). Such peaks could identify SNPs that affect splicing, so-called splicing QTLs, or other RNA processing steps.

For each peak, we compared the estimate of allelic imbalance for that putative isoform (peak) to the estimate in the associated gene containing that peak. Although these could be highly correlated, they might not be for biological reasons, such as differential transcription usage, or technical reasons, such as some peaks being noisier than others, leading to loss of power at the gene level, etc. To explore this, we compared each isoform peak’s estimated allelic imbalance levels to that for the associated gene and observed very high concordance, with many being significant in both (83.3% of genes with at least one significant peak were significant at the gene level; 25% of genes significant at the gene level had at least one significant peak in that gene), and with only 16.6% being significant at the peak level but not the gene level (Fig. S7c). The *KANSL1* gene stands out visually because it has 37 peaks, 28 of them tested for allelic imbalance, (once for each of two SNPs), with points for each of the two SNPs forming a vertical line (Fig. S7d).

The strong agreement between peak- and gene-level analysis provides evidence that the population-based phasing used to produce the phased VCF used by our pipeline produced reasonable results. If the phasing was low quality, we would expect there to be many peaks within the same gene with opposite directions of effect due to flips in the phasing between those peaks.

### Allelic imbalance versus eQTL analysis

Whether allelic imbalance provides more power than standard eQTL analysis has not, to our knowledge, been explored in the context of single-cell or single-nucleus RNA-seq data. As such, we performed a downsampling analysis of our non-selected PD data, running both eQTL analysis with pseudobulk followed by Matrix eQTL[56] (Methods) and allelic imbalance analysis with our beta-binomial pseudobulk method. We limited our analysis to the SNP/gene pairs identified previously in excitatory neurons from bulk data (Fig. 5d). In particular, we randomly downsampled to different numbers of individuals and counted the significant hits with eQTL analysis and allelic imbalance analysis (Fig. 5f). At all levels of downsampling, allelic imbalance analysis outperforms eQTL analysis with more significant hits, suggesting that allelic imbalance has more power. We acknowledge that it is unclear how much this result depends on the choice of statistic and normalization used for the eQTL analysis. We used Matrix eQTL due to its popularity, but it is possible that these results may change if we use an alternative statistical model, e.g., negative binomial instead of normal distribution, mixed model instead of pseudobulk, different covariates, etc., or apply an alternative normalization, e.g., quantile normalization instead of counts per million followed by ComBat.

### Differential allelic imbalance

Our work so far has focused on identifying SNPs and genes for which the percentage of reads assigned to one allele is significantly different from 50%. To extend such analyses, we explored methods to evaluate for each SNP whether the proportion of reads assigned to each allele of a gene is significantly different between different groups, e.g., glia *vs.* neurons, cases *vs.* controls, etc. We compared our pseudobulk and mixed model approaches along with the published DASESC_bb method[16] for their ability to identify allelic imbalance between the PD and healthy control samples with the PD SNP/gene pairs (Methods). Note we did not test the DAESC_mix method because it is not relevant to our study as it is used for gene-level analysis rather than SNP-level analysis. We also did not correct for any other confounders, although we recognize that unlike standard allelic imbalance analysis, differential allelic imbalance analysis can be confounded by individual-level covariates, such as age and sex, and correcting for confounders can be especially important when comparing between conditions. In terms of runtime, unsurprisingly the pseudobulk approach is by far the fastest (median of 2.25 seconds over all cell types), followed by our mixed model approach (median of 82.0 seconds over all cell types), followed by DAESC (median of 1772 seconds over all cell types) (Fig. S8a). In terms of performance, the effect sizes (Fig. S8b) and p values (Fig. S8c) estimated by all three methods showed very high agreement, particularly between the DAESC and glmmTMB mixed models, as expected. In terms of the number of SNP/gene pairs with significant imbalance, DAESC identified one, glmmTMB identified two, and pseudobulk identified five, although it is unclear whether these differences are meaningful. These results were obtained using default settings for each method, so that it is possible the results could be improved with optimization. For example, with DAESC_bb many of the genes did not reach the method’s convergence cutoffs (though were still reported), so that it might be possible to run the method for more iterations, though this would entail an even longer runtime for this method that is the slowest of the three.

Taken together, these analyses suggest that DAESC and the glmmTMB mixed model are, perhaps unsurprisingly, very comparable, as is pseudobulk to a slightly lesser extent, though glmmTMB is faster than DAESC, and pseudobulk is even quicker.

## Discussion

Allele-specific expression, and allelic imbalance more specifically, offers a powerful tool for understanding how genomic variation affects transcriptomic expression. Here we showed how choices in experimental design and computational analysis affect downstream results and can provide more power to detect allelic imbalance.

Our study shows that read length is an important consideration. Although long reads generated with ONT and PacBio sequencing detect more phased UMIs per read than short reads with Illumina sequencing (Fig. 3b), we favor the latter due to the lower cost per read leading to more power to detect ASE for the same cost. We acknowledge that this may change as sequencing technologies advance. Additionally in the future, long-read sequencing could be used to better understand isoform-level ASE — we showed that long reads could be used to quantify isoform-level ASE by assigning long reads to their isoform of origin and examining transcriptional regulation at a finer level of detail (Fig. S4). By contrast, short-read analysis had to use Sierra peaks (Fig. S4b, S7a-d) or similar methods, a less easily interpretable and less powerful measure of isoform-level ASE.

We also showed that hybrid selection of snRNA-seq libraries improved our power to detect allelic imbalance with no evidence of reference bias in the resulting libraries (Fig. 4). Moreover, the sequencing resources required are reduced when focused on the relevant regions of the transcriptome (Fig. 4c,d). This approach has exciting potential for following up GWAS loci to further elucidate the causal SNPs and affected genes – assigning function to variation.

In addition to these experimental advances, we provide tools for computational analysis to extend ASE in new directions. In particular, we show there is high agreement between mixed model and pseudobulk methods for detection of allelic imbalance, with slightly better false positive rate control in the pseudobulk approach (Fig. 5b,c). We also developed a novel approach that shows that allelic imbalance can be extended beyond the gene level (Fig. S7a-d), opening the possibility of isoform-level analysis even in droplet-based single-cell and single-nucleus transcriptomic data. All the computational methods developed for this study are open source, allowing others to build on them both in terms of extending the methodology and for analysis of new or existing datasets.

There are many possible extensions to the work we present here. Recent work[57] has suggested that it is possible to use single-cell ASE data to explore trans-eQTLs. Further, we did not consider fine mapping[58] or combining eQTL analysis with ASE information[29], both directions with great promise. On the biological side, we focused on the use of ASE to help understand inherited SNPs and small indels, but with slightly different methodologies one could also consider using it to explore *de novo* mutations, copy number variants (CNVs) and larger structural variations, somatic mutations in otherwise healthy cells, such as with clonal hematopoiesis of indeterminate potential (CHIP)[59, 60], or even somatic mutations in cancer[61]. We foresee the use of our ASE methods with spatial data[62], as our pipeline has built-in support for this already. Taken together, there are many possible ways for single-cell and single-nucleus ASE can be exploited to provide interesting biological insights in the future.

## Conclusions

The emergence of single-cell and single-nucleus RNA-seq technologies for human genomics has opened the possibility of a better understanding of human disease at the cell type and state level. The ASE methods developed in our study enable the extraction of greater insights into eQTLs and the effects of genome variation on cellular function. We demonstrate optimized experimental design and computational methods that enable more powerful future studies in this area.

## Methods

### Samples

We obtained three aliquots of ∼33 mg of Frontal Cortex (BA9) fresh-frozen tissue from each of the two GTEx samples. Fresh-frozen tissue samples from human middle temporal gyrus of controls, individuals with ILBD, and PD patients from the ASAP Parkinson Cell Atlas in 5D (PD5D) were obtained from the Arizona Study of Aging and Neurodegenerative Disorders/ Brain and Body Donation Program at Banner Sun Health Research Institute[63] and will be detailed in Lin et al. (manuscript in preparation; https://cloud.parkinsonsroadmap.org/collections/asap-parkinson-cell-atlas-in-5d-pd5d/overview, https://zenodo.org/doi/10.5281/zenodo.8384742).

### Nuclei isolation

We isolated nuclei from two frozen post-mortem human brain cortex tissues as previously described[64] with the following modifications. 1) We added 20 µl Recombinant RNase Inhibitor (Takara, 2313A) into 4 ml EZ prep lysis buffer (Sigma, NUC101) in the first incubation step and another 4 µl Recombinant RNase Inhibitor in the second incubation step. 2) We passed the lysate through a 40 µm cell strainer (VWR, 21008-949) after the tissue grinding. 3) We did the optional wash with Phosphate Buffered Saline (PBS; Invitrogen, AM9625) containing 0.01% BSA (Sigma, B6917) and 0.04 U/µl Recombinant RNase Inhibitor. 4) We filtered the nuclei through a 20 µm strainer (Sysmex, 04-004-2325) before the final counting using a Cellometer K2 Image Cytometer (Nexcelom Bioscience, CMT-K2-MX-150).

For the PD samples, we cryosectioned frozen temporal cortex tissue vertically. We transferred ten pieces of 50 µm thick sections to 2 pre-cooled pellet microtubes (five pieces each) in the cryostat. We used a pestle to homogenize the sample in each tube ten times with 1.0 ml Lysis buffer (prepared from 8 ml Nuclei Pure Prep (Sigma, NUC201-1KT), 8 μl 1 M DTT (Sigma, 43816), 80 μl 10% Triton X-100 (Sigma, T8787)) on ice. For each tube, we transferred 1.0 ml of homogenate to a 2.0 ml microcentrifuge tube and incubated on ice for 5 min, mixing once during the incubation. We then filtered the homogenate with a 70 μm strainer (Thermo Fisher Scientific, 2236548), centrifuged 500 x g at 4°C for 5 min, removed the supernatant, and incubated the pellet on ice for 5 min with 500 µl Nuclei wash and resuspension buffer (NWRB; prepared from 14.8 ml Dulbecco’s Phosphate-Buffered Saline (DPBS; Gibco, 14190-144), 150 µl BSA (Thermo Fisher Scientific, AM2616), 75 µl Recombinant RNase Inhibitor (40 U/ml; Takara, 2313B)) without resuspension. We then added an additional 1.0 ml NWRB to each tube and resuspended the pellet followed by centrifuging 500 x g at 4°C for 5 min. We resuspended the with 2 ml of NWRB with DAPI (prepared with 5.5 ml NWRB, 11 µl DAPI (Thermo Fisher Scientific, 62247; 5 mg/ml in water)) and filtered with a 40 μm flowmi cell strainer (Bel-art, H13680-0040). We sorted nuclei with a FACSAria III cell sorter (BD Biosciences). We isolated nuclei by fluorescence activated cells sorting based on DAPI and size, FSC and SSC (FSC at 5V Filter ND1, SSC at 180V Filter 488/10, DAPI by 405 Laser at 277V Filter 450/50) using a 100 μM nozzle and a Flow Cytometry Size Calibration Kit (Invitrogen, F13838).

### Illumina single-nucleus RNA-seq and hybrid selection

We loaded nuclei of each GTEx sample onto three channels, with each pair named as channel 1, 2, or 3 (Fig. S9) of a Chromium Single Cell 3’ Chip B (10x Genomics, 1000073) aiming to recover 10,000 cells per channel. We prepared the snRNA-seq libraries with the Chromium Single Cell 3’ Library & Gel Bead Kit v3 (10x Genomics, 1000075) following the manufacturer’s protocol. We pooled all six libraries based on molar concentrations and sequenced on a NextSeq 500 instrument (Illumina, SY-415-1001) using High-Output v2.5 Kit (75 cycles) flow cells (Illumina, 20024906) with 28 bases for read 1, 55 bases for read 2, and 8 bases for Index read 1.

We also performed hybrid selection with 750 ng of the remaining libraries from channel 2 of each sample, using the SureSelectXT reagent kit, HSQ (Agilent, G9611A) and a set of custom baits (see Hybrid selection bait design section) following the vendor’s protocol (SureSelectXT Target Enrichment System for Illumina Paired-End Multiplexed Sequencing Library Protocol, version C3) with the following modifications. 1) We added 1 µl of each universal blocking oligo, TS-p5 and TS-p7 (IDT, 1016184, 1016188), to each sample library in the denaturing step. 2) We amplified the captured libraries by mixing 50 µl of Amp Mix (10x Genomics, 220119), 2 µl each of 10 µM P5 (5’-AATGATACGGCGACCACCGA-3’) and P7 (5’-CAAGCAGAAGACGGCATACGA-3’) primers, and 30 µl of library captured beads in a 100 µl reaction volume with the following thermocycling conditions, 45 seconds at 98°C, 9 cycles of 15 seconds at 98°C and 30 seconds at 60°C and 30 seconds at 72°C, then 1 minute at 72°C before hold at 4°C. 3). We purified the amplified PCR product with 1.8x volumes of AMPureXP SPRI beads (Beckman Coulter, A63881). We pooled libraries based on molar concentrations and sequenced on a NextSeq 500 instrument (Illumina, SY-415-1001) using High-Output v2.5 Kit (75 cycles) flow cells (Illumina, 20024906) with 28 bases for read 1, 55 bases for read 2, and 8 bases for Index read 1.

We then pooled the original and targeted libraries at a 2:1 molar concentration and sequenced them together on a NovaSeq 6000 (Illumina, 20012850) with a S1 Reagent Kit v1.5 (200 cycles) flow cell (Illumina, 20028318) with 28 bases for read 1, 202 bases for read 2, and 8 bases for Index read 1.

For each PD sample, we used about 30,000 - 40,000 events (∼15,000 nuclei) per channel for 10x Chromium GEM generation and barcoding following the manufacturer’s protocol (Chromium Single Cell 3 GEM, Library & Gel Bead Kit v3, 10x Genomics, 1000075). We quantified the libraries using the KAPA Library Quantification Kit Illumina Platforms (Kapa BioSystems, KK4873) and sequenced them on a NextSeq 550 instrument (Illumina, SY-415-1002) using High-Output v2.5 Kit (150 cycles) flow cells (Illumina, 20024907) with 28 bases for read 1, 120 bases for read 2, 10 bases for Index read 1, and 10 bases for Index read 2.

We performed hybrid selection for PD snRNA-seq libraries with the same methods as for the GTEx samples with the following exceptions. 1) To obtain sufficient material for hybrid selection, we amplified each individual library with the Targeted Gene Expression Assay (10x Genomics, CG000293 Rev C) using the Library Amplification Kit (10x Genomics, 1000249) and started with 25 ng of each library for six cycles of PCR amplification. We purified the PCR products with 1.0x volumes of AMPureXP SPRI beads (Beckman Coulter, A63881) and freshly prepared 80% ethanol. We grouped the 85 libraries into 11 pools for hybrid selection. For sequencing, we further pooled the hybrid selection into 3 “super pools.” 2) The hybrid selected libraries were sequenced on a NovaSeq 6000 (Illumina, 20012850) with a S1 Reagent Kit v1.5 (200 cycles) flow cells (Illumina, 20028318) with 28 bases for read 1, 150 bases for read 2, 10 bases for Index read 1, and 10 bases for index read 2.

### PacBio RNA-seq

We constructed standard PacBio libraries with ∼230ng of the remaining cDNAs generated from channel 1 of each sample above using the SMRTbell Express Template Prep Kit 2.0 (Pacific Biosciences, 100-938-900) following the manufacturer’s protocol. We sequenced the libraries on the Sequel II platform (Pacific Biosciences, 101-447-000) with 2 SMRT cells for each library.

### MAS-seq

We prepared the first two MAS-seq libraries with 1 µl (∼7.5 ng) of cDNA from channel 1 of each sample for sequencing each library on one SMRT cell on the Sequel IIe platform (Pacific Biosciences, 101-986-400) according to the published protocol[42].

For the Revio sequencing, we processed the same two 10x Chromium 3’ snRNA-seq cDNA products using the MAS-Seq for 10x Single Cell 3’ Kit (Pacific Biosciences, 102-659-600, protocol Version 1.03) with the following modifications. To account for the limited input cDNA of 7 ng, we added an extra cycle to the TSO-PCR step. To increase library yield, we increased the MAS-PCR volume to 50 µl, the input purified cDNA to 80 ng, and decreased the PCR cycle number to 7. Following MAS PCR, the SMRTbell cleanup bead ratio was altered from 1.5x Bead-Addition-Volume-to-Reaction-Volume to 1.35x. Finally, the following steps: Post Array Ligation, Post DNA Damage Repair, and Final Cleanup were purified with a 1.55x SMRTbead cleanup. Following MAS library construction, libraries were quantified with a Qubit dsDNA High Sensitivity Assay kit (Thermo Fisher Scientific, Q32854) and a Femto-Pulse Genomic DNA 165 kb Kit (Agilent, FP-1002-0275). We sequenced each of the libraries on one SMRT cell on the Revio platform (Pacific Biosciences, 102-090-600).

### Oxford Nanopore Technologies RNA-seq with R2C2

We prepared Oxford Nanopore Technologies (ONT) libraries with the Rolling Circle Amplification to Concatemeric Consensus (R2C2) method[65] with the following modifications. First we re-amplified cDNA by combining 20 ng of the remaining cDNA from channel 2 of each sample together with 50 µl of NEBNext High-Fidelity 2x PCR Master Mix (New England Biolabs, M0541S), 2 µl of 12 µM PCR Primer1 (5’-CTACACGACGCTCTTCCGATCT-3’), 2 µl of 12 µM PCR Primer2 (5’-AAGCAGTGGTATCAACGCAGAGT-3’) in a 100 µl PCR reaction under the following conditions: 45 seconds at 98°C, 10 cycles of 10 seconds at 98°C and 15 seconds at 62°C and 3 minutes at 72°C, then 4 minutes at 72°C. We then purified the PCR products with 95 µl ProNex Size-Selective Purification System beads (Promega, NG2001) following the vendor’s protocol. We used 200 ng each of this reamplified cDNA in the circularization reaction. We made ONT libraries with 1000 ng each of the elongated cDNA with the Ligation Sequencing Kit (Oxford Nanopore Technologies, SQK-LSK109) following the vendor’s protocol and loaded each library on both a GridION X5 (Oxford Nanopore Technologies, GRD-X5B002) with a R9.4.1 flowcell (Oxford Nanopore Technologies, FLO-MIN106D) and a PromethION P48 (Oxford Nanopore Technologies, PRO-SEQ048) with a R9.4.1 flow cell (Oxford Nanopore Technologies, FLO-PRO002).

### PacBio sequence processing

We performed error correction for reads generated in CCS mode after transferring them from the Sequel II or on-board the Sequel IIe and Revio with the vendor’s ccs software (v5.0.0 for Sequel II, v.6.0.0 for Sequel IIe, v7.0.0 for Revio, https://github.com/PacificBiosciences/pbccs, RRID:SCR_021174) [66] and default settings (--all --suppress-reports --num-threads 232 --all-kinetics --subread-fallback). With these settings, reads failing error correction were automatically discarded and only passing reads were presented in a single BAM[67] file for downstream analysis. Each read was affixed with an auxiliary BAM tag rq indicating overall read quality ranging from 0.0 ≤ rq < 0.99 for corrected reads with predicted accuracy < Q20, and rq ≥ 0.99 for corrected reads with predicted accuracy ≥ Q20.

PacBio IsoSeq data was processed according to the vendor’s recommended workflow (https://isoseq.how/, v3.4.0, RRID:SCR_025481). Briefly, we removed IsoSeq adapters using PacBio’s lima tool (v2.6.0, RRID:SCR_025520) with options --guess 75 and --guess_min_count 1. We subsequently refined transcript reads to properly orient cDNA sequences, remove poly(A) tails, and retain only full-length non-chimeric (FLNC) transcripts using the isoseq3 refine command and default settings. Next, in preparation to error-correct transcripts, we clustered sequences using the isoseq3 cluster command and default settings. Finally, we collapsed sequences into a consensus sequence using the isoseq3 collapse command and default settings.

We processed the MAS-Seq sequencing HiFi reads into individual S-reads representing the original cDNA sequences using Skera (v1.2.0, https://skera.how/, RRID:SCR_025482) from the standard PacBio Toolkit. We processed this BAM file with the lima (v2.9.0) command, followed by the isoseq (v 4.0.0) tag command with --design T-12U-16B. This was followed by isoseq refine with the --require-polya tag, and then by isoseq correct with the associated barcode whitelist from Cell Ranger. This was transformed into a FASTA file with samtools fasta -T CB,XM for mapping downstream. Unlike the standard PacBio processing, duplicate sequences were not removed, as that was performed during the quantification step.

### ONT sequence processing

We performed basecalling for reads generated on Oxford Nanopore instruments (GridION and PromethION) on-board the respective instruments with the vendor’s Guppy software (v4.0.14; https://community.nanoporetech.com/downloads/guppy/release_notes, RRID:SCR_023196) and default settings. With these settings, reads failing basecalling were automatically discarded and only passing reads were presented in FASTQ files for downstream analysis.

### R2C2 concatemer segmentation, QC, and annotation

We applied C3Poa (git commit hash 0dab17c69930976c209c9a8f1df8ac037e570455, RRID:SCR_025484)[36] to split concatemers at provided splint sequences, error-correct resulting subreads using a partial order alignment approach, and discard reads that could not be properly processed (C3POA.py -r <fastq> -s <splint.fasta> -c /c3poa.config.txt -l 100 -d 500 -n 32 -g 1000 -o out).

To error-correct cell barcodes, we applied Starcode (v1.4, RRID:SCR_025483)[68] with default parameters to cluster all putative cell barcode sequences with entries from the 10x Genomics 3’ 3M-february-2018.txt CBC whitelist, subsequently replacing observed barcodes with cluster centroids.

### Long-read alignment and metrics calculation

We aligned reads to the GCA_000001405.15_GRCh38_no_alt_analysis_set.fna.gz (<u>Which</u> <u>human reference genome to use? (lh3.github.io)</u>) (herein referred to as “grch38_noalt”) reference sequence using appropriate platform-specific tools (Nanopore: minimap2 (v2.17-r941, RRID:SCR_018550)[69] -ayYL --MD --eqx -x splice -R <read group> -t <number of CPUs><grch38_noalt reference> <reads.fastq>; PacBio: pbmm2 (v1.4.0, https://github.com/PacificBiosciences/pbmm2, RRID:SCR_025549) align <input.bam> <ref.fa> <output.bam> --preset ISOSEQ --sample <sample_name> --strip --sort --unmapped; MAS-Seq: minimap2 (v2.28-r1209) --secondary=no -ayY --MD --eqx -x splice:hq --cs=long). The MD tag encoding mismatched and deleted reference bases was computed post-alignment using the SAMtools calmd command (v1.19.2, RRID:SCR_002105)[70]. We computed genome-wide and per-chromosome coverage metrics using MosDepth (v0.3.1, RRID:SCR_018929)[71], and read length/sequence identity metrics using NanoPlot (git commit hash e0028d85ec9e61f8c96bea240ffca65b713e3385, RRID:SCR_024128)[72] and custom scripts.

### Illumina snRNA-seq analysis (75 base read 2)

We loaded the Cell Ranger (v3.0.2, RRID:SCR_017344) output for the Illumina (75 base read 2) sequencing of the six 10x channels into Seurat (v3.1.1, RRID:SCR_016341)[73]. We removed cells with <500 genes detected. We normalized the data using the NormalizeData command, with a size factor of 1,000,000 and normalization.method=”LogNormalize” and identified variable genes with FindVariableFeatures with default arguments. We then scaled the data with the ScaleData command (with vars.to.regress=”nFeature_RNA”) followed by calculating principal component analysis (PCA) with the RunPCA command, generated a UMAP with Run UMAP (using dims=1:17), and performed clustering with the FindNeighbors command (with dims=1:17) followed by the FindClusters command (with resolution=2). To identify doublets, we used scrublet (v0.1, RRID:SCR_018098)[74]. We then labelled cells by cell type using known marker genes (*GAD1* for inhibitory neurons, *SLC17A7* for excitatory neurons, *SNAP25* for neurons in general, *PLP1* for oligodendrocytes, *PDGFRA* for oligodendrocyte precursor cells, *SLC1A3* for astrocytes, *CSF1R* for microglia, and *FLT1* for endothelial cells). These data were used for hybrid selection bait design.

### Illumina snRNA-seq (130 base read 2)

For clustering and cell type annotation of the GTEx Illumina data, we loaded expression information from the STARSolo output from both samples with the ReadInSTARSolo function from our scAlleleExpression package. We created a Seurat (v.4.0.0, RRID:SCR_016341) [75] object, filtering out cells with < 250 genes, and normalized the data with the NormalizeData function (normalization.method=”LogNormalize” and scale.factor=10000). We identified variable genes using FindVariableFeatures with default parameters, ran scGBM (v0.1.0, RRID:SCR_025518)[76] on the count data with 20 dims and the batch corresponding to sample of origin, and added the result as a dimensionality reduction object to the Seurat object. We performed clustering and UMAP projection on this with the FindNeighbors, FindClusters, and RunUMAP functions in Seurat with standard parameters, except with 20 dimensions as input and with the scGBM reduction instead of PCA. We added pointers to the output gene-level ASE counts and SNP information into the metadata using the LoadPipeline function in scAlleleExpression. We used scds(v1.6.0, RRID:SCR_021541)[77] on each sample separately to assign each cell a doublet score, remove clusters with high doublet scores, and then used known marker genes (see Illumina snRNA-seq analysis (75 base read 2 section)) to help identify cell types for each of the remaining clusters. We also removed cells with scds score > 1 as possible doublets.

For MAS-Seq data, we took the same approach, except the input count data was from UMI-tools (as described above) instead of STARSolo.

### Illumina snRNA-seq (PD-related samples)

We started from BCL files and processed them through the Cell Ranger (v5.0.1, RRID:SCR_017344) [78] suite of tools and cellranger mkfastq, cellranger counts (--include-introns) to demultiplex and map the data to the human genome using the human transcriptome reference obtained from Cell Ranger website’s download page (reference 2020-A (July 7, 2020), GRCh38, Ensembl 98)[79]. We subjected each sample to quality control prior to data aggregation. Cells with less than 200 unique genes or unique genes greater than three median average deviations above the median unique genes across all samples were discarded as low coverage or putative doublets, respectively. We also removed cells with greater than 5% of their reads mapping to the mitochondrial genome. In each sample, we only retained genes with a minimum count of one in at least three cells. We then processed the resulting aggregated matrix using a Seurat (v4.3.0, RRID:SCR_016341) workflow in R[75, 80]. We normalized the count data to give counts per ten thousand and these values + 1 were then natural log transformed. (NormalizeData, params: normalization.method = “LogNormalize”, scale.factor = 10000). We identified the 2000 most variable genes (FindVariableFeatures, params: selection.method = “vst”) and scaled the data (ScaleData) across all genes. We then used the 2000 most variable features for dimensionality reduction using glmPCA (0.2.0, https://github.com/willtownes/glmpca, RRID:SCR_025517) (RunGLMPCA)[81]. We processed the data with Harmony (v1.2.0, RRID:SCR_022206)[82] to account for covariates, correcting the glmPCA for the snRNA-seq batch, sample, age bracket (binned into 10-year intervals), RIN bracket, PMI bracket, and sex, with theta being set at 0.4 for each covariate. We used an elbow plot to help determine the ideal number of harmony dimensions (60) to pass to the FindNeighbors function, which was followed by cluster identification using the Leiden algorithm, in which a range of resolutions was tested (0.5 – 1.6) to identify a suitable value (FindClusters, params: resolution = 1.5, algorithm = 4, method = “igraph”)[83]. We then ran UMAP (RunUMAP) to visualize the resulting clusters[84]. We removed clusters with less than 200 cells or representing less than 20 samples as part of cluster quality control. We identified doublet cells using the scDblFinder package (v1.12.0, https://plger.github.io/scDblFinder/, RRID:SCR_022700), by running the function scDblFinder on a per sample basis. All doublet cells were removed as well as any clusters that consisted of 30% or more doublet cells. We subsequently re-ran the UMAP to clean up the visualization.

### Hybrid selection bait design

To select 100 genes to target in the GTEx snRNA-seq data, we used all six channels from the 75 base read 2 data and collected QC metrics (Supplementary Table 5) for all genes in the reference (original (pre-2020) GRCh38 reference from 10x Genomics). These metrics included the percentage of nuclei expressing these genes in each cell type and overall, their average expression (average logTPM) in each cell type and overall, the length of the gene (extracted from the GTF file in the Cell Ranger reference), the log fold change in each cell type relative to others, percentage and number of UMIs phased in each GTEx sample. We then performed these filtering steps:

1. **Basic filtering:** We required genes to be expressed in >1% of cells (overall), with >1% phased UMIs in both GTEx samples.
2. **Cell Type Specific:** We focused on three cell types, microglia, astrocytes, and excitatory neurons. For each cell type, we examined all the genes that were upregulated (FDR adjusted p-value < 0.05, logFC > 0) in that cell type relative to others based on MAST (v1.16.0, RRID:SCR_016340)[85] and selected 10 of the genes with a mix of genes with high and low expression in the associated cell type. To do this, we ordered the genes by the percentage cells of that cell type expressing that gene and picked the genes at equal intervals, ensuring we got some expressed in a high percentage and some in a low percentage. Similarly, we also picked another 10 genes among these cell type specific genes to have a mix of lengths. We ordered the genes by length and picked the genes at equal intervals. This gave us 60 selected genes.
3. **eQTL specific:** We also looked at known eQTLs from bulk RNA-seq of cortex from GTEx[31] with (a) SNPs that were heterozygous in one of our two samples, (b) genes that were expressed in >10% of cells in the dataset, and (c) SNPs located in either an intron or exon. We ordered the genes by the effect size of their biggest eQTL, and as above picked genes at equal intervals, picking 20 of them. This gave us a mix of genes whose eQTLs have stronger and weaker effect sizes. This gave us 20 more selected genes.
4. **Percentage UMI Phased:** Starting from genes that were expressed in >10% of cells in the dataset, we ordered the genes by the percentage of UMIs that were phased for that gene and picked 10 genes at equal intervals, giving us a mix of genes with higher and lower percentage of phased UMIs. This gave us 10 more selected genes.
5. **Overall Expression:** Finally, we ordered the genes by their average expression and picked genes at equal intervals, giving us genes of mixed expression levels. This gave us 10 more genes, for a total of 100 selected genes (Supplementary Table 5).

For these 100 genes, we identified all the SNPs that were present in at least one of those genes using bedtools (v2.29, RRID:SCR_006646) intersect[86] and that were heterozygous in at least one of our samples. We further limited the number of SNPs by choosing those that had >1 read overlapping them in sample 1 and >1 read in sample 2. We then produced the final targeted regions by choosing a region with 130 bp on either side of each SNP and merging overlapping regions. Finally, we used these regions as input to the Agilent SureSelect design tool with overnight digestion, 2x coverage, with boosting, and stringent repeat masking. The bait regions are listed in Supplementary Table 6.

For the 101 genes we targeted in our PD dataset, listed in Supplementary Table 5, we used an initial version of SNP/gene pairs chosen based on eQTL analysis of PD GWAS loci using snRNA-seq data from the human MTG samples (Parker, Scherzer et al., manuscript in preparation).

For these genes, we used a similar approach to select regions to target as above. We processed each non-selected sample from the PD dataset one by one using a Nextflow (DSL 1.0, RRID:SCR_024135)[87] pipeline. First, we processed each sample with the same version of the pipeline used for GTEx samples with the same reference. Next, we used sinto (v0.7.2.2, https://github.com/timoast/sinto, RRID:SCR_025498) to extract all reads overlapping the genes of interest from the output BAM file. We transformed the resulting BAM file into a BED file with the bedtools bamtobed command (with the -split flag) and extracted entries overlapping heterozygous SNPs in that sample with bedtools intersect. We used an awk command (awk ‘{if($6==“+”){print $1“\t”$2“\t”($3+200)}if($6==“-”){print $1“\t”($2-200)“\t”$3}}’) to extend these BED files by 200 bp in the 3’ direction. With bedtools sort, we sorted the BED file and combined overlapping entries in the extended BED file with bedtools merge. We combined the resulting BED files from all samples with the cat command and sorted them with bedtools sort. We processed this BED file with a Python script that read each line in one by one and extracted regions covered by at least 10 entries in the BED file, saving them to a BED file of targeted regions. Finally, we ran it through the Agilent SureSelect design tool with overnight digestion, 2x coverage, with boosting, and stringent repeat masking. The bait regions are listed in Supplementary Table 6.

### Read processing pipeline

To extract ASE information from 10x Chromium RNA-seq data, we built the single-cell allele-specific expression processing pipeline, which is a Nextflow pipeline (DSL 1.0)[87] (v1, https://zenodo.org/records/10696967). The pipeline takes in FASTQ files, a barcode whitelist (built in whitelists exist for Visium, 10x Chromium v2, and 10x Chromium v3), a VCF with phased genotype data, and reference files (including a STAR reference and a GTF). These files were preprocessed, then passed to STARSolo (v2.7.8, RRID:SCR_021542) with arguments described in https://github.com/seanken/ASE_pipeline/blob/main/QuantPipeline.no.conda.nf_line103 (using default parameters for the Nextflow pipeline for all analyses unless otherwise stated). We used a STAR reference built using the GTF and FASTA from the original (pre-2020) GRCh38 reference from 10x Genomics. In particular, we used the WASP[29, 30] capability built into STAR (v2.7.8) to help mitigate the effects of reference bias. For the analysis without intronic reads, we used the same pipeline except with Gene as the argument for –soloFeatures instead of GeneFull (used in all other analyses). The pipeline then used a htsjdk (v2.21.1, RRID:SCR_024036)[67] based Java application to calculate cell-level ASE from the STARSolo BAM file. For each read in the file, the application extracted information about the read from that the SAM tags. Multimapping reads and reads that did not pass the WASP filter were excluded. For each heterozygous SNP overlapping the read, the allele (alternative *vs.* reference) was extracted from the vA tag and the read’s allele was assigned based on this information. If SNPs within the same read disagreed on the allele to be assigned, the read was assigned to the more common allele if it accounted for >95% of SNPs in the read, else the read was excluded. The UMI, CBC, and gene for the remaining reads were extracted from the associated tags, and information about the UMI/CBC/gene/allele were added to a HashMap. This HashMap was then used to assign UMI/CBC/gene triplets to allele (reference, alternative, or ambiguous, where ambiguous corresponds to UMI/CBC/Gene triplets that are assigned to different alleles by different reads) and the results were saved to an output file. Although we used this pipeline for extracting ASE information from our GTEx sample datasets and for generating regions to target for the PD data, a newer version of the pipeline (also at https://zenodo.org/records/10696967), which was updated to use Nextflow DSL (v2.0), STARSolo (v2.7.10) and allow BAM files as input, but otherwise identical functionality was used for the PD datasets in all figures, with a different reference GTF but the same FASTA (GTF downloaded from https://ftp.ensembl.org/pub/release-98/gtf/homo_sapiens/Homo_sapiens.GRCh38.98.gtf.gz).

For long-read data, we wrote a Java application (https://zenodo.org/uploads/12531178) that takes in a BAM file annotated with cell barcodes, UMIs, and genes of origin, as well as a phased VCF. We annotated the gene of origin with featureCounts (v2.0.1, RRID:SCR_012919)[88] (with tags -L -t gene -g gene_name -o gene_assigned -s 2 -R CORE) and extracted expression information with UMI-tools (v1.1.2, RRID:SCR_017048)[89]. The computational processing approach for this tool is the same as that for the short-read data, except instead of extracting SNP information from tags in the BAM file for each read, we loaded the genotype information into a HashMap and used that to identify which locations in each read overlapped a heterozygous SNP. We used the same GTF file used for the other GTEx samples.

For downsampling analysis of Illumina data, we downsampled BAM files from our pipeline using Picard Tools (v2.27.5, http://broadinstitute.github.io/picard, RRID:SCR_006525) DownsampleSam before processing them with our jar file for counting ASE. Similarly, for the read length analysis, we trimmed reads to different lengths with the command “awk -v cutlen=$cutlen ‘{if(NR%2==0){print substr($0,0,cutlen)}else{print $0}}’” (where cutlen is a variable containing the length to trim to) before feeding them into the Illumina pipeline. For downsampling the long-read data, we used Picard Tools DownsampleSam to downsample the BAM files before running the above long-read pipeline.

In addition to the ASE pipeline, we also ran Cell Ranger[78] count (v3.0.2, RRID:SCR_017344) on each Illumina sample, with default parameters, except for the expected number of cells which were determined by the number of cells loaded in each 10x Chromium channel. This was performed on the Cell Ranger GRCh38 reference with introns added.

### Downstream analysis package

We built an R package, scAlleleExpression (v1, https://zenodo.org/records/10697254), for downstream analysis. The package had two purposes. First, it was built to load data from our read processing pipeline into R, a functionality which was used to load data for all ASE-based analysis in this manuscript. In particular, the package was built around a central data frame with one row per cell, with each column corresponding to either metadata, or serving as a pointer to a particular pipeline output. This metadata table was used to load ASE information at the SNP-level and gene-level expression information (as calculated by STARSolo). This was achieved by loading the gene-level ASE for each sample and the phased genotype at the SNPs of interest. Samples with non-heterozygous genotypes were excluded, and the ASE of the remaining samples in the gene of interest were aligned so that the allele with the reference value of the SNP was set as the reference allele of that gene. This enabled the use of reads both directly overlapping a SNP and those not directly overlapping it and was used by default for SNP-level analysis in our manuscript. Alternatively, it was possible to directly load the SNP-level ASE values, which are based only on the reads directly overlapping the given SNP. We used this last functionality to load data for all analyses involving genotype data without phasing information, as well as for analyses looking at reference bias. All analysis in the paper was performed with scAlleleExpression v1.

Second, the package also had statistical tools built in to enable allelic imbalance analysis. In particular, once one loaded ASE information (as described above), there were two main approaches to testing for allelic imbalance in the presence of multiple individuals. These statistical tests were used for that statistical analysis of the PD data. In the first approach using pseudobulk data, it was possible to fit a beta-binomial regression model with the aod package (v1.3.2, https://cran.r-project.org/package=aod, <u>10.32614/CRAN.package.aod</u>, RRID:SCR_025516) and extract the p-values and effect sizes for the coefficient of interest (more often than not the intercept term). Alternatively, if one was using single-cell level data, one could fit a beta-binomial mixed model (with a random effect corresponding to sample of origin) with the glmmTMB package (v1.1.5, RRID:SCR_025512)[52] and extract the p-values and effect sizes for the coefficient of interest (most often the intercept term). For gene-level allelic imbalance within a given sample, which we used for the statistical analysis of the GTEx data in this manuscript, the package can be used to fit a beta-binomial regression model with the aod package on cell-level ASE data and extract the p-values and effect sizes for the coefficient of interest (most often the intercept term).

We also built a Python package, scASE_py (v1, https://zenodo.org/records/11582140), to load the output of the pipeline into Python as a pandas (v1.0.5, RRID:SCR_018214)[90] data frame. This package also provides an interface to the scDali package. This package was not directly used for the data analysis in this manuscript.

### Differential allelic imbalance analysis

To test for differential allelic imbalance with the glmmTMB and pseudobulk methods, we used the same approach as for standard differential analysis (see above), except with the formula ∼Condition, where Condition corresponded to case *vs.* control status.

For DAESC (v0.1.0, https://zenodo.org/records/8329900)[16], we filtered the SNP/gene pairs as for the pseudobulk and mixed model approaches. For each SNP/gene pair, we used the cell-level ASE information as input for DAESC_bb with the settings niter=200, niter_laplace=2, num.nodes=3, optim.method=“BFGS”, and converge_tol=1e-8, and a model matrix built model.matrix(∼Conditon,data=meta), where meta was the cell-level meta data. We then collected the results and reported them back.

### Calling SNPs from snRNA-seq data

To call SNPs from snRNA-seq data, we ran cellsnp-lite (v1.2.3, RRID:SCR_025515)[91] to call the genotype of the SNPs from the GTEx VCF (using the -R flag), using the BAM file and filtered cell list from STARSolo, and the flags -p 8 –minMAF 0.05 –minCOUNT 10 –gzip – genotype. We took the output cellSNP.base.vcf.gz VCF file and used Python to create a VCF with those that were likely heterozygous SNPs (those with DP > 10 and the AD/DP ratio between 0.1 and 0.9). We then added a column of genotype information to the resulting VCF (all 0|1 genotypes since we chose the heterozygous sites) and used the resulting VCF for downstream analysis.

### Extracting read information for long-read data

To extract read length information from our long-read data, we used pysam (v0.15.3, https://github.com/pysam-developers/pysam, RRID:SCR_021017) to load each read one by one. We loaded the CIGAR information with cigartuples and counted the number of mapped bp.

To extract information about the number of indels and SNVs in the long- and short-read data, we used pysam to load each read one by one. The CIGAR string was extracted as above. The CIGAR string was then processed to count the number of matched bp, substitutions, insertions, and deletions.

### Sierra analysis

We used Sierra (v0.99.24, RRID:SCR_025510)[41] to better understand allelic imbalance at a level closer to that of isoforms. For each sample, we processed the BAM files and filtered cell lists produced by STARSolo through a Nextflow (DSL 1.0) pipeline built to extract Sierra peaks. This pipeline indexed the BAM file, made junction files with the RegTools (v0.5.2, https://zenodo.org/records/7521875)[92] junction command with the -s 1 flag, then called peaks with the FindPeaks command in Sierra. These peaks were then merged with the MergePeakCoordinates command and annotated with AnnotatePeaksFromGTF, using BSgenome.Hsapiens.UCSC.hg38::Bsgenome.Hsapiens.UCSC.hg38 as the genome reference. We modified this annotation by (1) not reporting the polyA and polyT categories and (2) including 5’ UTR and 3’ UTR as exon.

To count the phased UMIs in each peak, we used a Nextflow pipeline (DSL 1.0) that started by transforming the merged peaks into a BED file. Each entry in the BED file corresponded to a contiguous region in a Sierra peak, where BED entries coming from the same peak were given the same name. We annotated the STARSolo BAM file with these peaks using the command bedtools tag -names -s -tag PK -I $input -files $peaksBed where input was the BAM file from STARSolo and peaksBed was the BED file produced from the merged peaks. This added a new tag, PK, to each read in the BAM file with information about any Sierra peaks the read overlapped. Finally, we used the same jar file used for extracting gene-level ASE to extract Sierra-level ASE, except using the PK tag, which contained peak-level information, instead of the GN tag from the BAM file. The list of Sierra peaks for both datasets is in Supplementary Table 7.

### Isoform-level expression and ASE from long-read data

To extract isoform-level information from long-read data, we modified the gene-level long-read ASE pipeline – creating the genome mapping based isoform ASE pipeline. In particular, instead of using featureCounts for annotating reads with genes of origin, we built a custom Python script as part of our long-read processing pipeline (v1, zenodo.org/uploads/12531178) that annotated each read with the isoforms it overlaps. To do this, the script first loaded the GTF file into Python, creating two dictionaries. The first dictionary had one interval tree, which was built with intervaltree (v3.1.0, https://github.com/chaimleib/intervaltree, RRID:SCR_025514RRID), per chromosome, where the interval tree contains one interval per isoform from that chromosome (the interval consisting of the region covered by the isoform), including one such interval per unspliced gene. The second dictionary mapped from isoform names (the transcript id appended to the gene id) to a list of the regions in the genome covered by each exon. Once these dictionaries were constructed, each alignment was loaded in one at a time with pysam. A dictionary was created mapping each read to the maximum alignment score over all alignments. The alignments were then read through a second time. Those alignments that did not have the maximum alignment score for the associated read were excluded, as were reads with multiple alignments with the same maximum score. A list of all isoforms overlapping that alignment were extracted from the interval trees built from the GTF file using the overlap command. The regions covered by the exons of each isoform were extracted from the second dictionary and trimmed to the region covered by the read of interest. Each read was then transformed into a list of regions covered by that read with the get_blocks command, merging blocks that were within 10 bp of each other. Each of the isoform-level lists of exons were compared to the list from the read. We excluded isoforms where the overlap was < 95% of the length of the region covered by that isoform and < 95% of the region covered by the read. We then found the remaining isoform with the largest overlap. If there were other isoforms whose overlap was < 10bp shorter than this largest overlapping isoform, the read was discarded; otherwise, the read was assigned to the isoform with the largest overlap. The script generated a new BAM file containing the reads that were assigned to an isoform, with the isoform name being included in the XT tag.

For short-read data, we generated a modified version of the standard short-read pipeline by adding two additional steps. In the first step the genome-aligned BAM file from STARSolo (produced in the single cell pipeline) was annotated by isoform using the same script as for the long-read data, adding the results as a new tag (XT). The second step used the ASE quantification script from the short-read pipeline to quantify ASE at the gene level to instead quantify ASE at the isoform level, feeding in the isoform annotated BAM and using the XT tag instead of the standard gene tag, but otherwise the using the same settings as for gene-level analysis.

### Extracting cell-level read mapping information

For Illumina data, we calculated cell-level QCs with the CellLevel_QC tool (https://github.com/seanken/CellLevel_QC, https://zenodo.org/records/12794612). In particular, we annotated reads from STARSolo BAM files as exon, intron, or intergenic with bedtools tag -s -tag RE -i $bam -files $genes $exons $anti -labels N E A, where $genes, $exons, and $anti were GTF files produced by extracting the gene-level, exon-level, and antisense (gene-level with strand flipped) entries from the input GTF file, with the information stored in the RE tag. We used the resulting BAM file as input for the CellLevel_QC jar file to extract cell-level mapping information. In addition, we also annotated the BAM file from our pipeline with bedtools tag -i $bam -s -f 1 -files $bed1 $bed2 $bed3 $bed4 $bed5 -labels UTR3 UTR5 Exon Gene Anti -tag GR, where $bed1, $bed2, $bed3, $bed4, and $bed5 were GTF files produced by extracting the 3’ UTR, 5’ UTR, exon, gene, and antisense (gene with strand flipped) entries from the input GTF file, with the information stored in the GR tag. We processed this BAM file with a Python script to extract this information. We loaded reads into Python one by one with pysam and excluded multimappers (NH tag not equal to 1). We extracted the GR tag and used it to assign reads to different categories (see Fig. 2b). The script then counted, for each category, the percentage of reads that had a vW tag, which was the percentage overlapping a heterozygous SNP.

### eQTL analysis

For each sample in the PD dataset, we calculated the pseudobulk count data with a modified version of the LoadExpression function in our scAlleleExpression package, modified to perform pseudobulking while loading each sample using the rowSums command. We normalized the data to counts per million (CPM) and removed genes with low expression (those with less than 1 CPM expression in at least 75% of samples). We further normalized the data with ComBat (RRID:SCR_010974) [93] in the sva package (v3.38.0, RRID:SCR_012836) [94] using the batch information. We extracted the expression of the genes being tested and saved this information as a tsv file.

We calculated the genotype principal components (PCs) as follows. We converted the VCF to a GDS file with the snpgdsVCF2GDS command in the SNPRelate package (v1.24.0, RRID:SCR_022719)[95]. We used the same package for LD pruning using the command snpgdsLDpruning with autosome.only=TRUE, maf=0.05, missing.rate=0.05, method=“corr”, slide.max.bp=10e6, and ld.threshold=sqrt(0.1), and calculated PCA with snpgdsPCA. We then combined the top 5 PCs with the metadata from the pseudobulk step (namely age and sex) and saved them as a tsv file. We used the genotypeToSnpMatrix command in the VariantAnnotation package (v1.36.0, RRID:SCR_000074)[96] to convert the VCF into a genotype matrix, loading 100,000 SNPs at a time, excluding those with minor allele frequency < 0.05 in our dataset. We downsampled this file to the SNPs we wanted to test and saved the results as a SNP data file.

Finally, we ran Matrix_eQTL_engine from the MatrixEQTL package (v2.3, RRID:SCR_025513)[56] on the files we produced above, using useModel = modelLINEAR, pvOutputThreshold = 1, errorCovariance = numeric(), min.pv.by.genesnp = FALSE, and noFDRsaveMemory = FALSE. We extracted the eQTL results, filtered out those from SNP/gene pairs we were not testing, calculated the FDR, and saved the results.

For the downsampling analysis, we loaded the relevant matrices (expression, SNP, and meta data) into R with the read.table command, downsampled to the appropriate number of individuals, and then converted to a SlicedData object to be used with Matrix_eQTL_engine as above. In addition to saving the eQTL output, we saved the names of samples selected in each downsampling step to allow us to run allelic imbalance analysis on the exact same set of samples.

## Supplementary Information

### Supplementary Figures

**Fig. S1.**
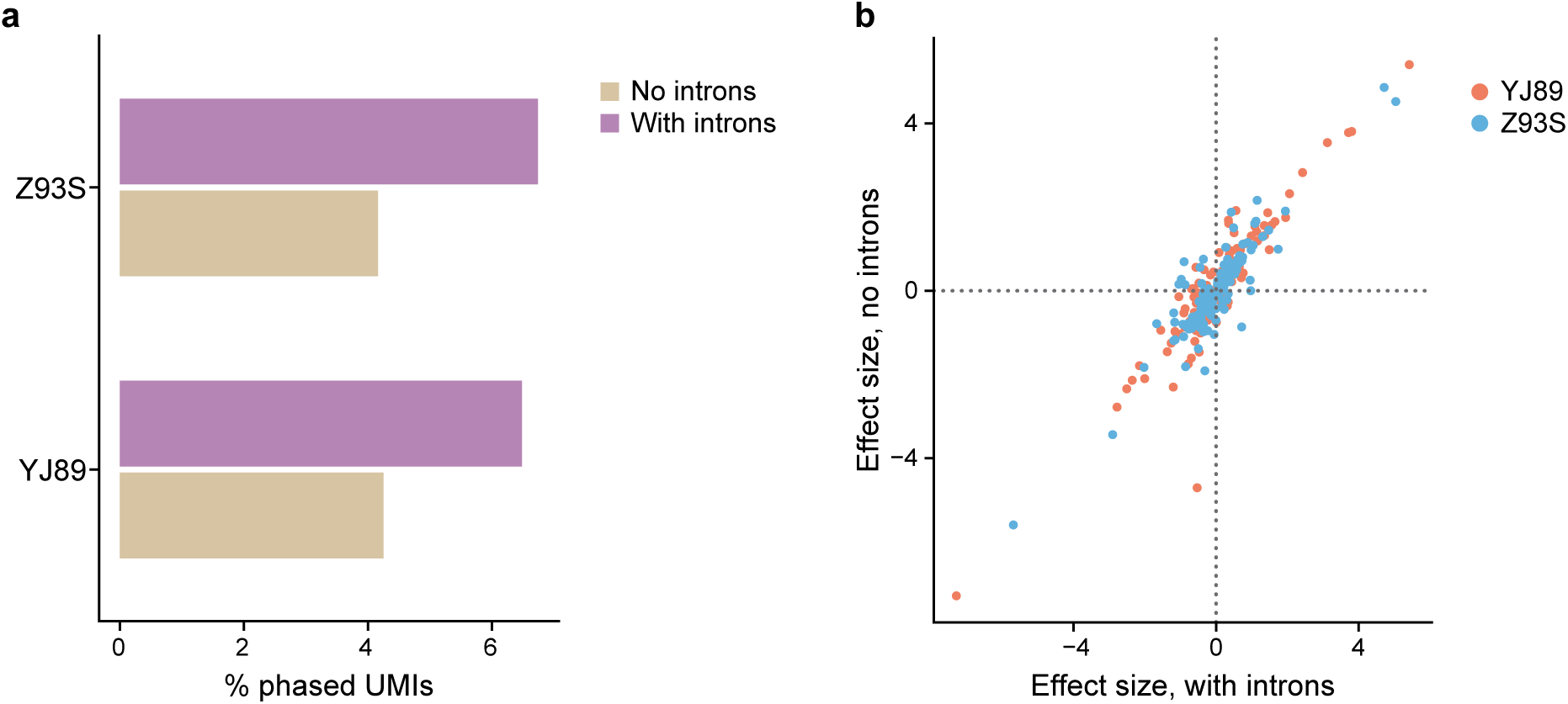
ASE with or without intron-aligned UMIs. **a** Comparison of the percentage of UMIs that are phased in each sample either including or excluding introns. **b** Comparison of the estimated allelic imbalance effect size for genes that are significant in either analysis with or without introns for each sample.

**Fig. S2.**
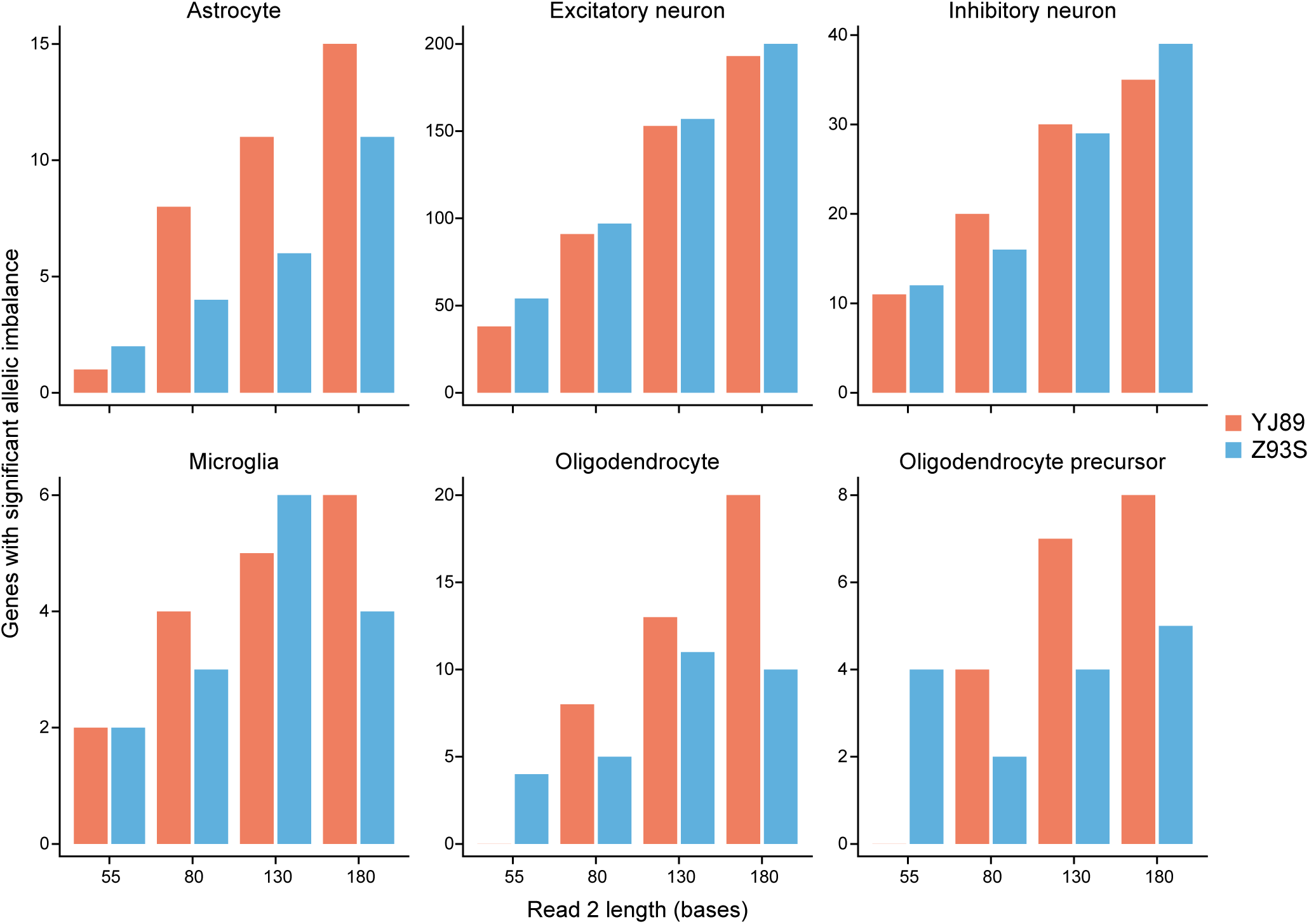
Read length effect on power in each cell type. Shown for each cell type is the number of genes with significant allelic imbalance for each sample at different read lengths.

**Fig. S3.**
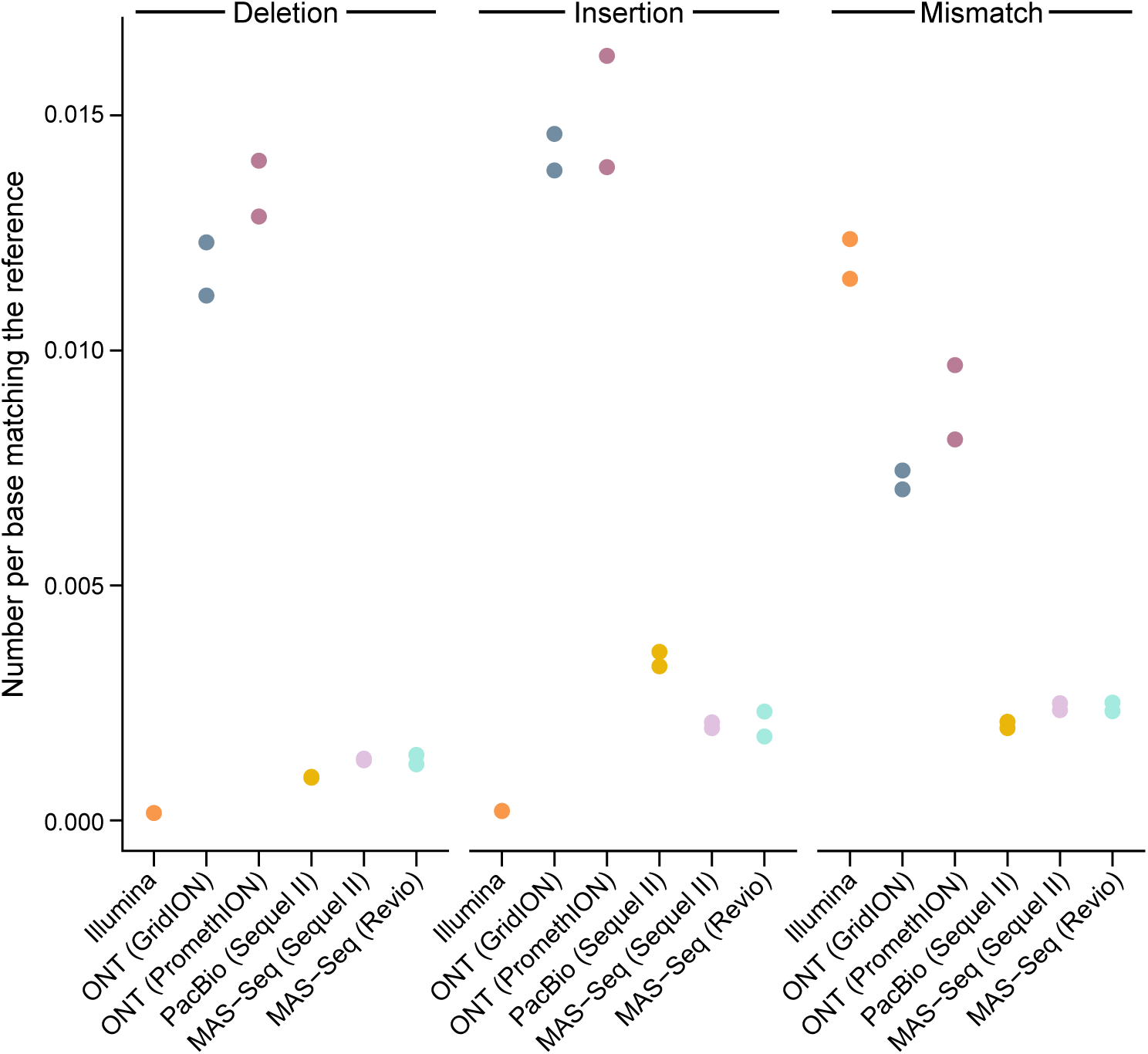
Long-read error rates. For each sequencing technology, shown is the frequency a base disagreed with the reference genome, with either an insertion, deletion, or mismatch. We counted matches (positions were the read matches the reference genome), mismatches (positions where the read maps to the genome but the nucleotide does not match the reference), insertions (locations with an insertion in the read relative to the reference genome), and deletions (locations with a deletion in the read relative to the reference genome, excluding splicing events). We then calculated the number of insertions, deletions, and mismatches divided by the number of matches and plotted the results.

**Fig. S4.**
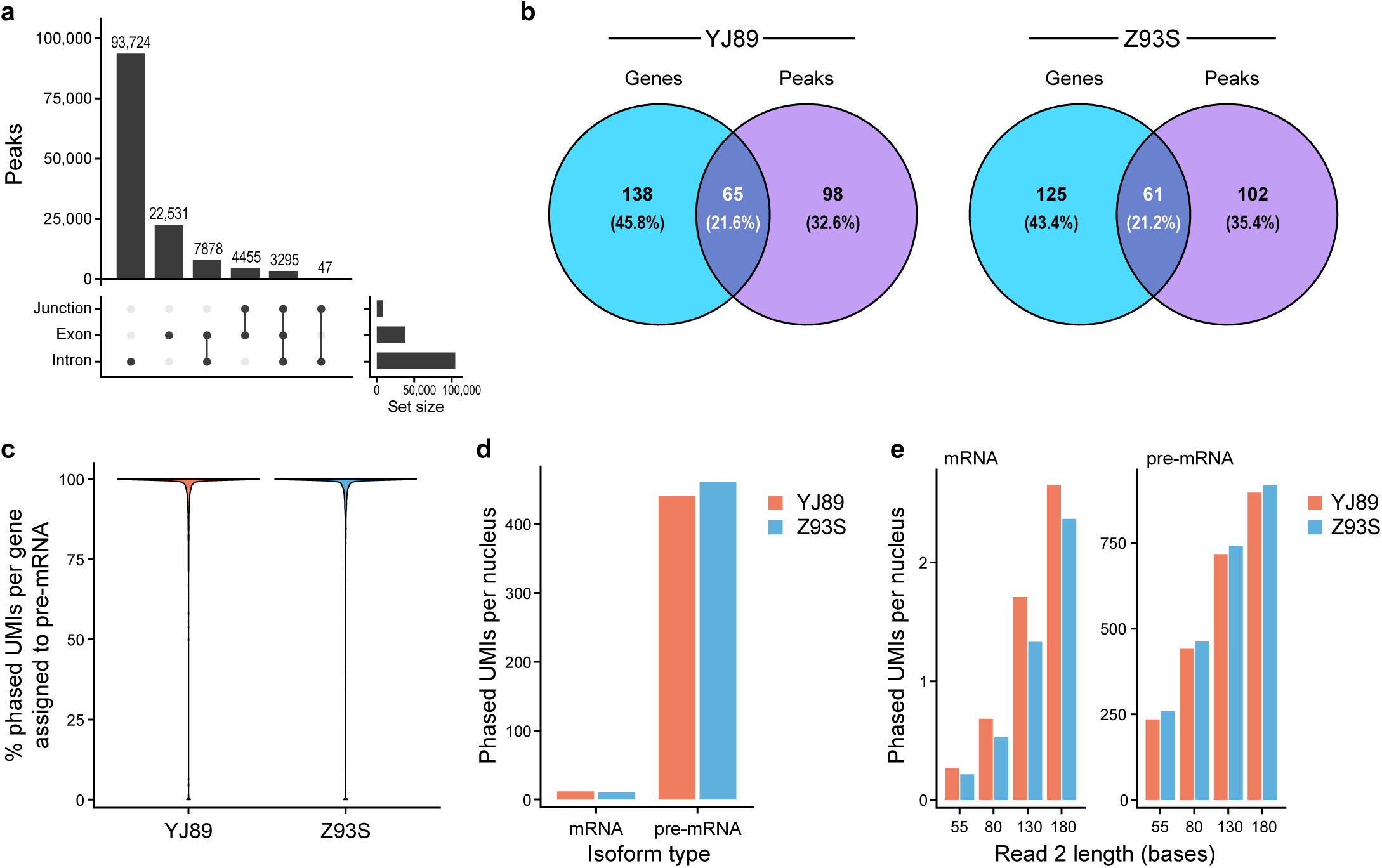
Isoform-level analysis. **a** UpSet plot of the number of peaks assigned to a genomic region according to Sierra’s peak annotation. Exon includes 5’ UTR and 3’ UTR peaks. **b** Venn diagram for each sample showing overlap between genes with significant allelic imbalance and genes with at least one Sierra peak with significant allelic imbalance. **c** Violin plot showing the percentage of phased UMIs coming from the unspliced pre-mRNA transcript for each gene with at least 10 phased UMIs in the isoform level analysis for each sample. **d** Phased UMIs per cell recovered with isoform-level analysis of the MAS-Seq data for both spliced and unspliced reads. **e** Phased UMIs per cell recovered with isoform-level analysis of the short-read data for spliced and unspliced reads with varying lengths of read 2.

**Fig. S5.**
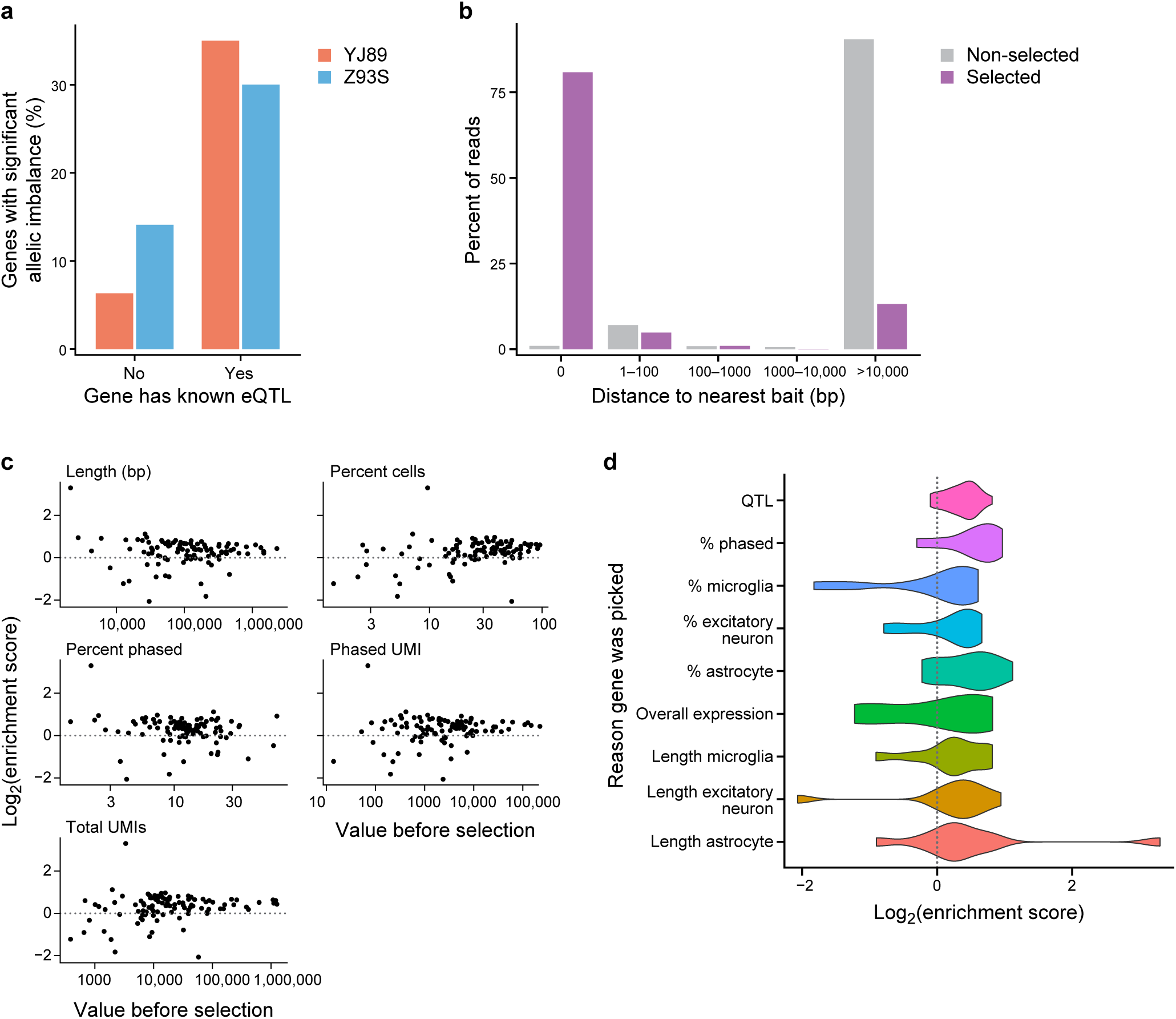
Hybrid selection metrics. **a** Percentage of targeted genes with significant allelic imbalance shown for genes with or without a known eQTL. **b** Distance of uniquely mapped reads from hybrid selection baits with and without selection**. c** Comparison of the log_2_ of the enrichment score (phased UMIs after selection / phased UMIs before selection) for each selected gene to different metrics for each gene without selection. Total UMI: total UMIs assigned to that gene, Phased UMI: total phased UMIs assigned to that gene, Length: length of the gene in bp, Percentage Phased: percentage of Phased UMI divided by Total UMI, and Percentage Cells: percentage of cells expressing that gene. **d** Violin plots of the enrichment score for genes chosen by each criterion (Methods).

**Fig. S6.**
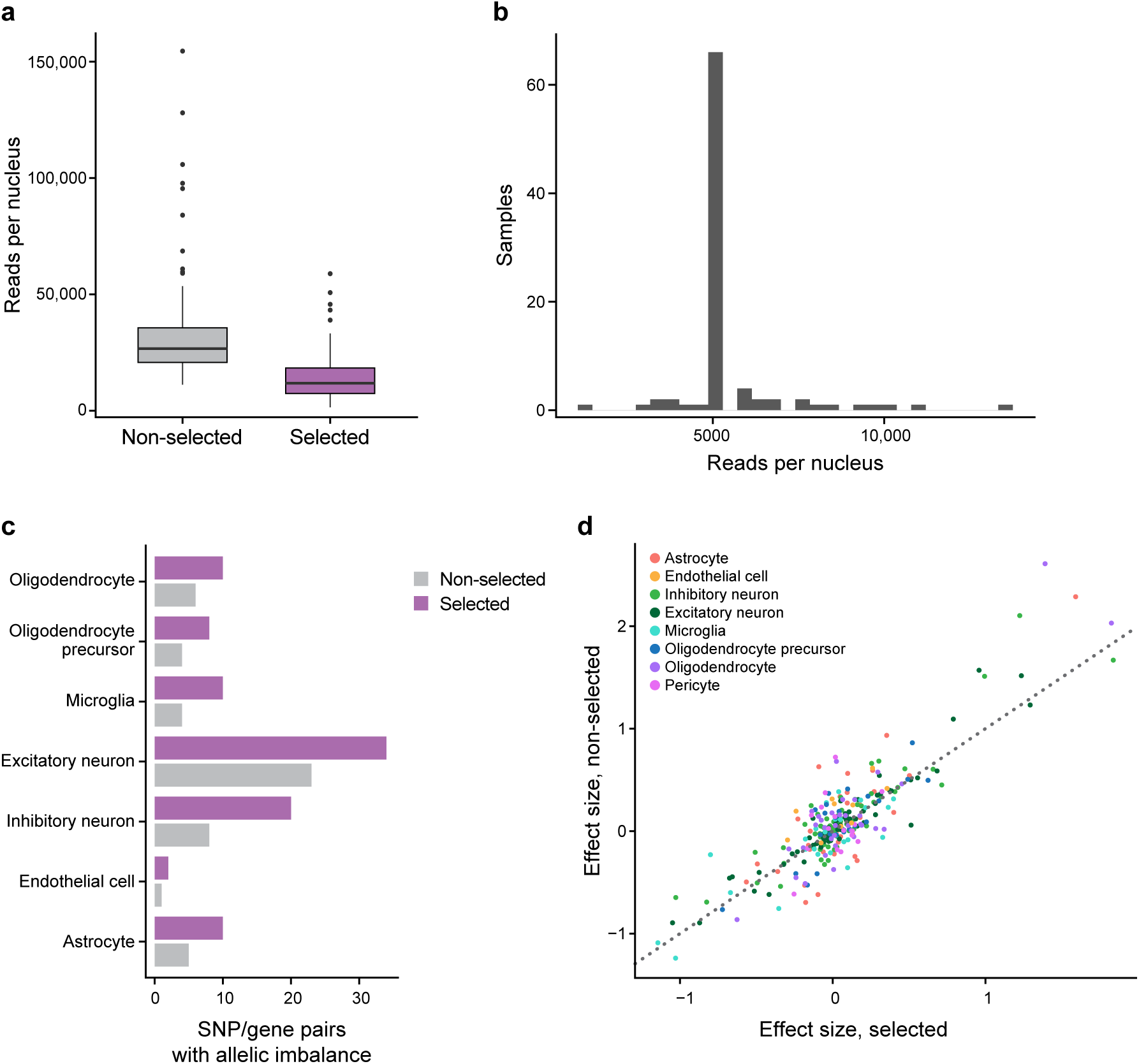
Additional hybrid selection analysis. **a** Boxplots of sequence coverage without selection and with selection. **b** Histogram of sequence coverage with downsampling. Overall, there are an average of 5,365 reads per cell in this dataset. Because the downsampling was based on the number of nuclei reported by Cell Ranger and these results were based on the number of nuclei in the final Seurat object (a smaller number of nuclei), some samples had more than 5,000 reads per nucleus. Those with fewer than 5,000 reads per nucleus were not downsampled. **c** Comparison of the number of SNP/gene pairs with significant allelic imbalance in each cell type for downsampled selected data and non-downsampled non-selected data. **d** Comparison of effect size estimates for each targeted gene and each cell type in the data with and without selection.

**Fig. S7.**
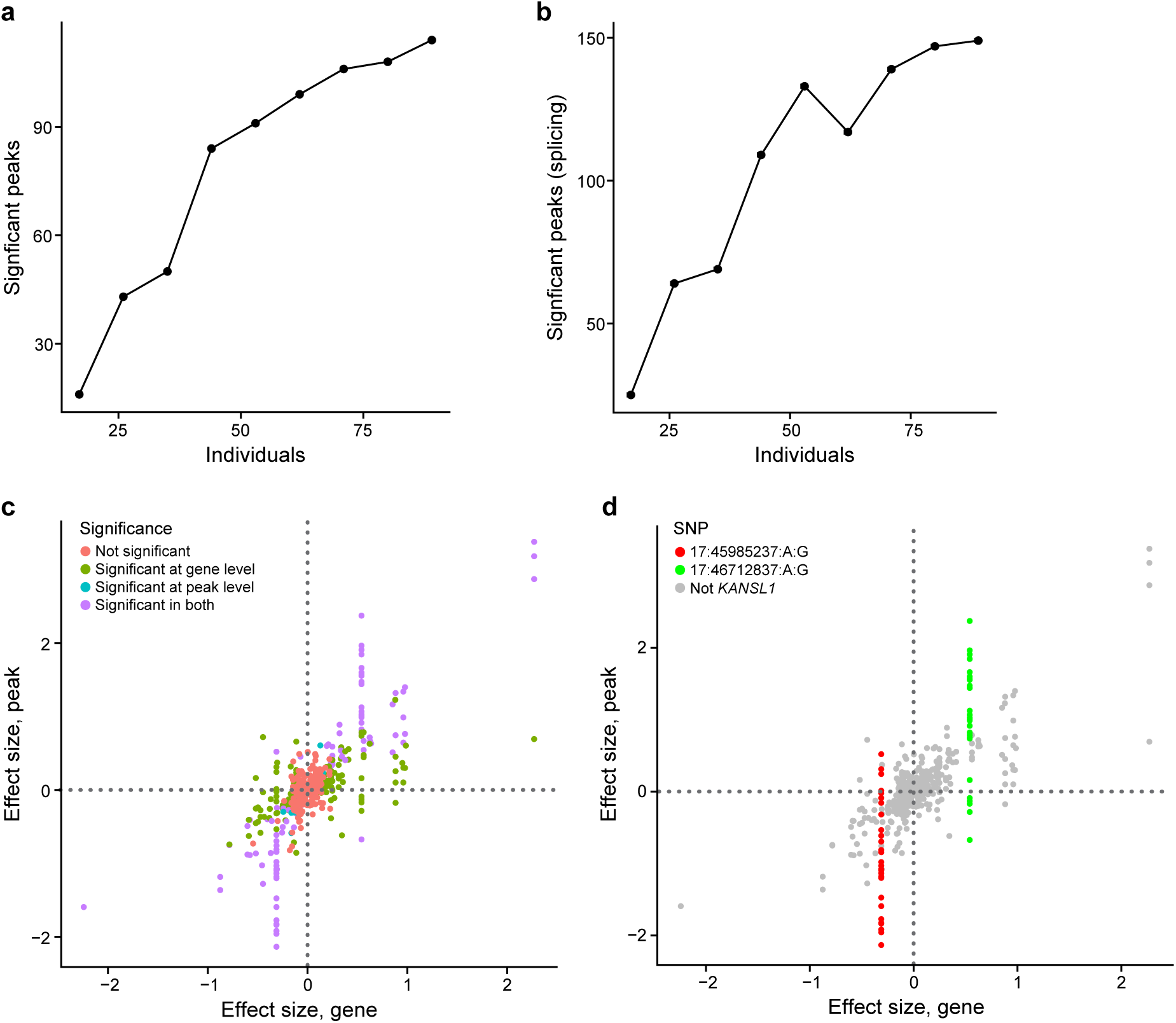
Sierra peak analysis. Effects of downsampling the number of individuals on Sierra peaks with significant allelic imbalance (**a**) and Sierra peaks with significant allelic imbalance relative to the other peaks in the same genes (**b**). **c** Estimated allelic imbalance for each peak *vs.* estimated allelic imbalance for the associated gene. **d** As in **c**, highlighting *KANSL1* peaks.

**Fig. S8.**
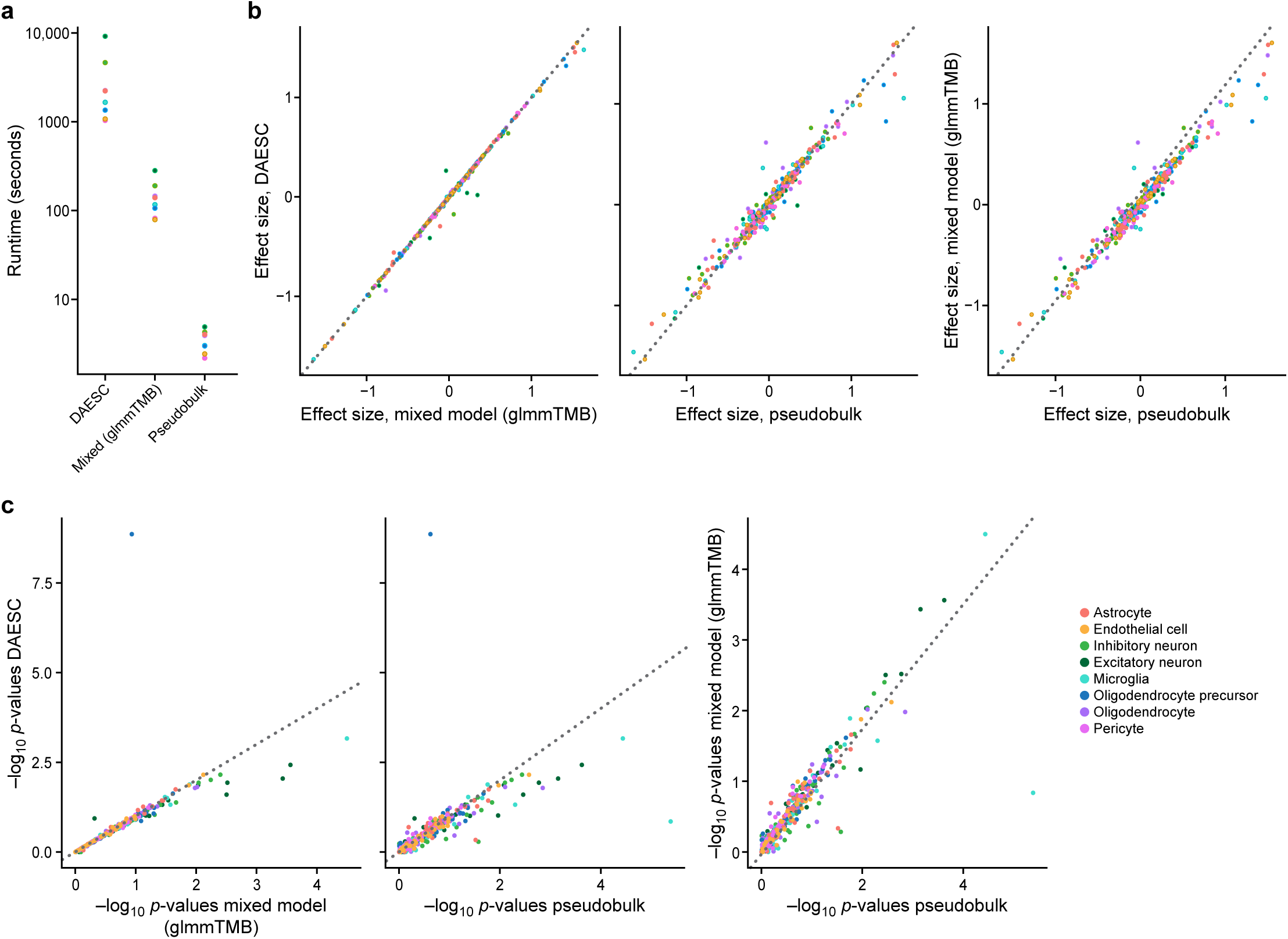
Differential allelic imbalance. **a** Runtime of each method for each cell type. Scatter plots comparing the estimated effect size (**b**) and the negative log_10_ p-values (**c**) from differential allelic imbalance analysis (PD *vs.* healthy controls) for each method in each cell type. All analyses used the PD data with selection and no downsampling.

**Fig. S9.**
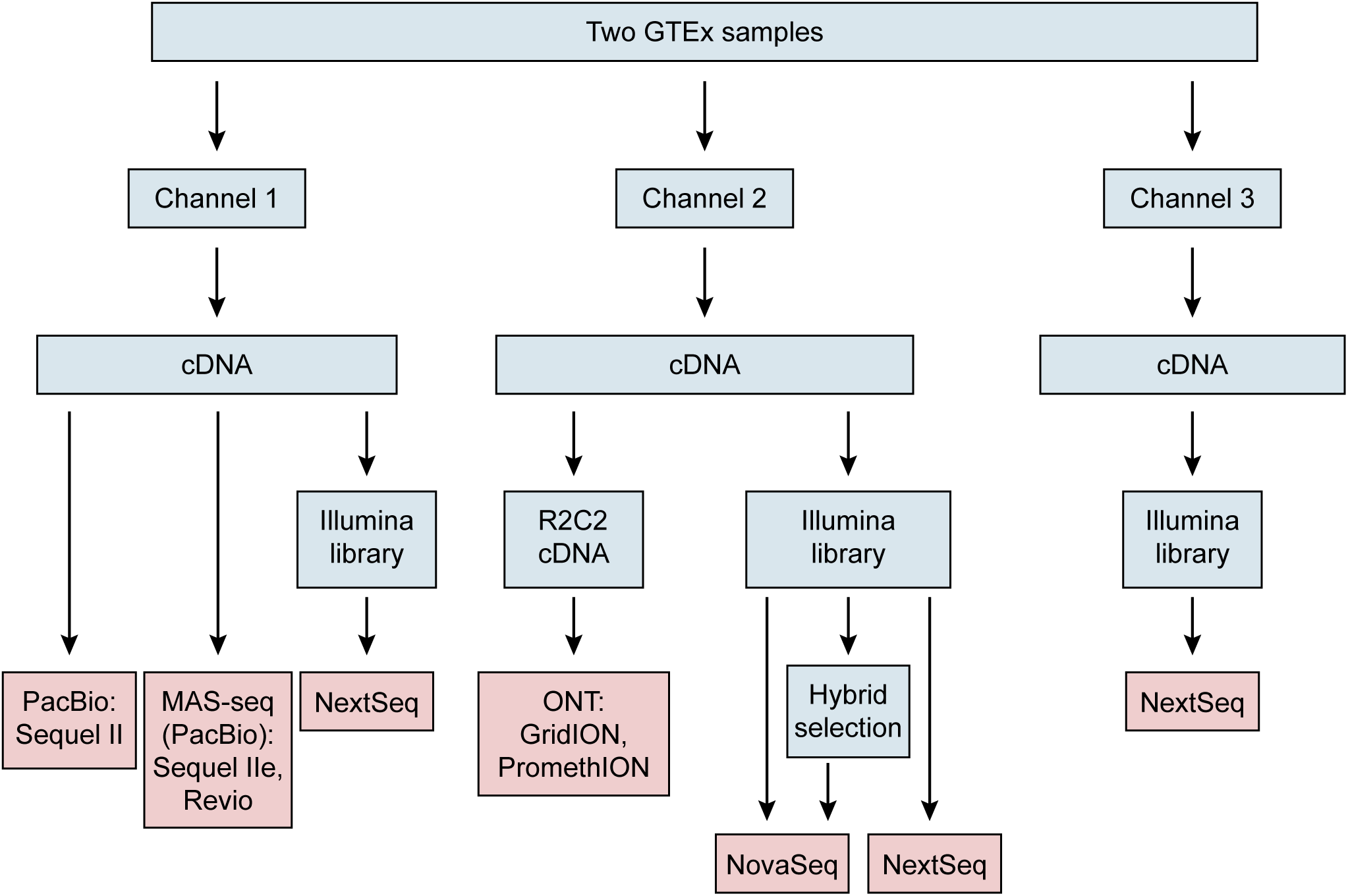
Experimental design. Schematic showing processing of each library with the GTEx samples.

### Supplementary Tables

**Supplementary Table 1** QC metrics reported by STARSolo and the ASE pipeline for GTEx samples

**Supplementary Table 2** Peak- and gene-level allelic imbalance results with GTEx data

**Supplementary Table 3** List of GTEx samples for long- and short-read data

**Supplementary Table 4** Allelic imbalance results for PD data with and without selection

**Supplementary Table 5** Selected genes in PD and GTEx data with gene-level QC metrics

**Supplementary Table 6** Targeted regions in selection experiments for PD and GTEx data

**Supplementary Table 7** Annotated Sierra peaks for PD and GTEx data

**Supplementary Table 8** Key Resource Table

## Declarations

### Ethics approval and consent to participate

Ethics approval and consent information for GTEx samples have been previously published[97]. For PD-related samples, brain autopsies were approved by the local IRB of the Banner Brain and Body Donation program. The PD-related studies described here were approved by the Institutional Review Boards of Mass General Brigham and Yale.

### Consent for publication

Not applicable

### Availability of data and materials

The snRNA-seq FASTQ files for the GTEx samples is available from dbGaP with the ascension phs003749.v1.p1. For the Parkinson’s disease cohort, raw data are available in the ASAP Collaborative Research Network (CRN) Cloud in the “ASAP Parkinson Cell Atlas in 5D (PD5D)” dataset (DOI: 10.5281/zenodo.11585274, see the dataset “team-scherzer-pmdbs-sn-rnaseq-mtg”). The Parkinson’s data are also available in the ASAP CRN Cloud (DOI: 10.5281/zenodo.14270014, see the dataset “team-scherzer-pmdbs-sn-rnaseq-mtg-hybsel”). The annotation and counts for the GTEx data used in Figure 2 are available in the Broad Single Cell Portal under the accession SCP2873. See the Key Resource Table (Supplementary Table 8) for additional information.

Code from this study is available through a central GitHub repository, https://github.com/seanken/ASE_Central. This repository has links to the different repositories hosting the tools from this paper. The code from the pipelines and packages developed as part of this manuscript is also available through Zenodo; see the Key Resource Table (Supplementary Table 8) for specifics.

### Authors contributions

S.K.S., J.Z.L, A.R. and O.R.R. conceived the strategies used in this study. X.A. performed the snRNA-seq experiments with GTEx samples. S.K.S. developed the code and performed the ASE analysis. A.M.A. and A.S. performed the MAS-seq experiments. N.H. performed the hybrid selection experiments with PD samples with guidance from J.Z.L. M.G., J.T.S., and K.G. performed the long-read sequence processing. K.J. and K.G. assisted S.K.S. in analysis for selecting genes for hybrid selection of GTEx samples. Z. Liao and I.T. generated PD snRNA-seq data with guidance from C.R.S. C. B.-A. helped optimize FACS for PD single nuclei isolation J.P. and Z. Lin analyzed these PD data with guidance from X.D. and C.R.S. D.E.K. and Y.K. provided project management support for PD experiments. G.E.S. and T.G.B. provided PD brain samples. S.K.S., J.Z.L., X.A., A.S, K.G., A.A., and N.H. wrote the manuscript with input from the other authors.

## Acknowledgements

For the purpose of open access, the author has applied a CC BY public copyright license to all Author Accepted Manuscripts arising from this submission. We thank Beatrice Weykopf for help with the Key Resource Table, the Broad Genomics Platform for sequencing, Bryce van de Geijn for advice on building the computational pipeline, Leslie Gaffney for help with illustrations and figures, and Houlin Yu for sharing code that we used as a basis for our MAS-seq pipeline. We also acknowledge Kristin Ardlie and the GTEx project team for providing samples and genotype data. The GTEx Project was supported by the Common Fund of the Office of the Director of the National Institutes of Health, and by NCI, NHGRI, NHLBI, NIDA, NIMH, and NINDS. We are grateful to the Arizona Study of Aging and Neurodegenerative Disorders/ Brain and Body Donation Program at Banner Sun Health Research Institute, Sun City, Arizona for the provision of human biological materials. The Brain and Body Donation Program has been supported by the National Institute of Neurological Disorders and Stroke (U24 NS072026 National Brain and Tissue Resource for Parkinson’s Disease and Related Disorders), the National Institute on Aging (P30 AG019610 and P30AG072980, Arizona Alzheimer’s Disease Center), the Arizona Department of Health Services (contract 211002, Arizona Alzheimer’s Research Center), the Arizona Biomedical Research Commission (contracts 4001, 0011, 05-901 and 1001 to the Arizona Parkinson’s Disease Consortium) and the Michael J. Fox Foundation for Parkinson’s Research.

## References

1. Oelen R, de Vries DH, Brugge H, Gordon MG, Vochteloo M, single-cell e Qc, Consortium B, Ye CJ, Westra HJ, Franke L, van der Wijst MGP: Single-cell RNA-sequencing of peripheral blood mononuclear cells reveals widespread, context-specific gene expression regulation upon pathogenic exposure. Nat Commun 2022, 13:3267.

2. Perez RK, Gordon MG, Subramaniam M, Kim MC, Hartoularos GC, Targ S, Sun Y, Ogorodnikov A, Bueno R, Lu A, et al: Single-cell RNA-seq reveals cell type-specific molecular and genetic associations to lupus. Science 2022, 376:eabf1970.

3. Yazar S, Alquicira-Hernandez J, Wing K, Senabouth A, Gordon MG, Andersen S, Lu Q, Rowson A, Taylor TRP, Clarke L, et al: Single-cell eQTL mapping identifies cell type-specific genetic control of autoimmune disease. Science 2022, 376:eabf3041.

4. Ding R, Wang Q, Gong L, Zhang T, Zou X, Xiong K, Liao Q, Plass M, Li L: scQTLbase: an integrated human single-cell eQTL database. Nucleic Acids Res 2024, 52:D1010–D1017.

5. Kang JB, Raveane A, Nathan A, Soranzo N, Raychaudhuri S: Methods and Insights from Single-Cell Expression Quantitative Trait Loci. Annu Rev Genomics Hum Genet 2023, 24:277–303.

6. Castel SE, Aguet F, Mohammadi P, Consortium GT, Ardlie KG, Lappalainen T: A vast resource of allelic expression data spanning human tissues. Genome Biol 2020, 21:234.

7. Zou J, Hormozdiari F, Jew B, Castel SE, Lappalainen T, Ernst J, Sul JH, Eskin E: Leveraging allelic imbalance to refine fine-mapping for eQTL studies. PLoS Genet 2019, 15:e1008481.

8. Mohammadi P, Castel SE, Brown AA, Lappalainen T: Quantifying the regulatory effect size of cis-acting genetic variation using allelic fold change. Genome Res 2017, 27:1872–1884.

9. Almlof JC, Lundmark P, Lundmark A, Ge B, Maouche S, Goring HH, Liljedahl U, Enstrom C, Brocheton J, Proust C, et al: Powerful identification of cis-regulatory SNPs in human primary monocytes using allele-specific gene expression. PLoS One 2012, 7:e52260.

10. Guelfi S, D’Sa K, Botia JA, Vandrovcova J, Reynolds RH, Zhang D, Trabzuni D, Collado-Torres L, Thomason A, Quijada Leyton P, et al: Regulatory sites for splicing in human basal ganglia are enriched for disease-relevant information. Nat Commun 2020, 11:1041.

11. Larsson AJM, Johnsson P, Hagemann-Jensen M, Hartmanis L, Faridani OR, Reinius B, Segerstolpe A, Rivera CM, Ren B, Sandberg R: Genomic encoding of transcriptional burst kinetics. Nature 2019, 565:251–254.

12. Cuomo ASE, Seaton DD, McCarthy DJ, Martinez I, Bonder MJ, Garcia-Bernardo J, Amatya S, Madrigal P, Isaacson A, Buettner F, et al: Single-cell RNA-sequencing of differentiating iPS cells reveals dynamic genetic effects on gene expression. Nat Commun 2020, 11:810.

13. Heinen T, Secchia S, Reddington JP, Zhao B, Furlong EEM, Stegle O: scDALI: modeling allelic heterogeneity in single cells reveals context-specific genetic regulation. Genome Biol 2022, 23:8.

14. Mu W, Sarkar H, Srivastava A, Choi K, Patro R, Love MI: Airpart: interpretable statistical models for analyzing allelic imbalance in single-cell datasets. Bioinformatics 2022, 38:2773–2780.

15. Qi G, Battle A: Computational methods for allele-specific expression in single cells. Trends Genet 2024.

16. Qi G, Strober BJ, Popp JM, Keener R, Ji H, Battle A: Single-cell allele-specific expression analysis reveals dynamic and cell-type-specific regulatory effects. Nat Commun 2023, 14:6317.

17. Booeshaghi AS, Gao F, Pachter L: Assessing the multimodal tradeoff. bioRxiv 2023:2021.2012.2008.471788.

18. M Pn, Liu H, Bousounis P, Spurr L, Alomran N, Ibeawuchi H, Sein J, Reece-Stremtan D, Horvath A: Estimating the Allele-Specific Expression of SNVs From 10x Genomics Single-Cell RNA-Sequencing Data. Genes (Basel) 2020, 11.

19. Habib N, Avraham-Davidi I, Basu A, Burks T, Shekhar K, Hofree M, Choudhury SR, Aguet F, Gelfand E, Ardlie K, et al: Massively parallel single-nucleus RNA-seq with DroNc-seq. Nat Methods 2017, 14:955–958.

20. Bakken TE, Hodge RD, Miller JA, Yao Z, Nguyen TN, Aevermann B, Barkan E, Bertagnolli D, Casper T, Dee N, et al: Single-nucleus and single-cell transcriptomes compared in matched cortical cell types. PLoS One 2018, 13:e0209648.

21. Eraslan G, Drokhlyansky E, Anand S, Fiskin E, Subramanian A, Slyper M, Wang J, Van Wittenberghe N, Rouhana JM, Waldman J, et al: Single-nucleus cross-tissue molecular reference maps toward understanding disease gene function. Science 2022, 376:eabl4290.

22. Gaublomme JT, Li B, McCabe C, Knecht A, Yang Y, Drokhlyansky E, Van Wittenberghe N, Waldman J, Dionne D, Nguyen L, et al: Nuclei multiplexing with barcoded antibodies for single-nucleus genomics. Nat Commun 2019, 10:2907.

23. Wu H, Kirita Y, Donnelly EL, Humphreys BD: Advantages of Single-Nucleus over Single-Cell RNA Sequencing of Adult Kidney: Rare Cell Types and Novel Cell States Revealed in Fibrosis. J Am Soc Nephrol 2019, 30:23–32.

24. Hu P, Liu J, Zhao J, Wilkins BJ, Lupino K, Wu H, Pei L: Single-nucleus transcriptomic survey of cell diversity and functional maturation in postnatal mammalian hearts. Genes Dev 2018, 32:1344–1357.

25. Deng Q, Ramskold D, Reinius B, Sandberg R: Single-cell RNA-seq reveals dynamic, random monoallelic gene expression in mammalian cells. Science 2014, 343:193–196.

26. Hagemann-Jensen M, Ziegenhain C, Chen P, Ramskold D, Hendriks GJ, Larsson AJM, Faridani OR, Sandberg R: Single-cell RNA counting at allele and isoform resolution using Smart-seq3. Nat Biotechnol 2020, 38:708–714.

27. Wilson PC, Muto Y, Wu H, Karihaloo A, Waikar SS, Humphreys BD: Multimodal single cell sequencing implicates chromatin accessibility and genetic background in diabetic kidney disease progression. Nat Commun 2022, 13:5253.

28. Benjamin K, Dinar Y, Alexander D: STARsolo: accurate, fast and versatile mapping/quantification of single-cell and single-nucleus RNA-seq data. bioRxiv 2021:2021.2005.2005.442755.

29. van de Geijn B, McVicker G, Gilad Y, Pritchard JK: WASP: allele-specific software for robust molecular quantitative trait locus discovery. Nat Methods 2015, 12:1061–1063.

30. Asiimwe R, Dobin A: STAR+WASP reduces reference bias in the allele-specific mapping of RNA-seq reads. bioRxiv 2024:2024.2001.2021.576391.

31. Consortium G: The GTEx Consortium atlas of genetic regulatory effects across human tissues. Science 2020, 369:1318–1330.

32. Castel SE, Levy-Moonshine A, Mohammadi P, Banks E, Lappalainen T: Tools and best practices for data processing in allelic expression analysis. Genome Biol 2015, 16:195.

33. Choi Y, Chan AP, Kirkness E, Telenti A, Schork NJ: Comparison of phasing strategies for whole human genomes. PLoS Genet 2018, 14:e1007308.

34. Loh PR, Danecek P, Palamara PF, Fuchsberger C, Y AR, H KF, Schoenherr S, Forer L, McCarthy S, Abecasis GR, et al: Reference-based phasing using the Haplotype Reference Consortium panel. Nat Genet 2016, 48:1443–1448.

35. Edsgard D, Reinius B, Sandberg R: scphaser: haplotype inference using single-cell RNA-seq data. Bioinformatics 2016, 32:3038–3040.

36. Volden R, Palmer T, Byrne A, Cole C, Schmitz RJ, Green RE, Vollmers C: Improving nanopore read accuracy with the R2C2 method enables the sequencing of highly multiplexed full-length single-cell cDNA. Proc Natl Acad Sci U S A 2018, 115:9726–9731.

37. Stoler N, Nekrutenko A: Sequencing error profiles of Illumina sequencing instruments. NAR Genom Bioinform 2021, 3:lqab019.

38. LaPierre N, Pimentel H: Accounting for Isoform Expression in eQTL Mapping Substantially Increases Power. bioRxiv 2023:2023.2006.2028.546921.

39. He D, Gao Y, Chan SS, Quintana-Parrilla N, Patro R: Forseti: A mechanistic and predictive model of the splicing status of scRNA-seq reads. bioRxiv 2024:2024.2002.2001.577813.

40. Zhou R, Xiao X, He P, Zhao Y, Xu M, Zheng X, Yang R, Chen S, Zhou L, Zhang D, et al: SCAPE: a mixture model revealing single-cell polyadenylation diversity and cellular dynamics during cell differentiation and reprogramming. Nucleic Acids Res 2022, 50:e66.

41. Patrick R, Humphreys DT, Janbandhu V, Oshlack A, Ho JWK, Harvey RP, Lo KK: Sierra: discovery of differential transcript usage from polyA-captured single-cell RNA-seq data. Genome Biol 2020, 21:167.

42. Al’Khafaji AM, Smith JT, Garimella KV, Babadi M, Popic V, Sade-Feldman M, Gatzen M, Sarkizova S, Schwartz MA, Blaum EM, et al: High-throughput RNA isoform sequencing using programmed cDNA concatenation. Nat Biotechnol 2023.

43. Zee A, Deng DZQ, Adams M, Schimke KD, Corbett-Detig R, Russell SL, Zhang X, Schmitz RJ, Vollmers C: Sequencing Illumina libraries at high accuracy on the ONT MinION using R2C2. Genome Res 2022, 32:2092–2106.

44. Joglekar A, Hu W, Zhang B, Narykov O, Diekhans M, Marrocco J, Balacco J, Ndhlovu LC, Milner TA, Fedrigo O, et al: Single-cell long-read sequencing-based mapping reveals specialized splicing patterns in developing and adult mouse and human brain. Nat Neurosci 2024, 27:1051–1063.

45. Foox J, Tighe SW, Nicolet CM, Zook JM, Byrska-Bishop M, Clarke WE, Khayat MM, Mahmoud M, Laaguiby PK, Herbert ZT, et al: Performance assessment of DNA sequencing platforms in the ABRF Next-Generation Sequencing Study. Nat Biotechnol 2021, 39:1129–1140.

46. Zolotarov G, Grau-Bové X, Sebé-Pedrós A: GeneExt: a gene model extension tool for enhanced single-cell RNA-seq analysis. bioRxiv 2023:2023.2012.2005.570120.

47. Tardaguila M, de la Fuente L, Marti C, Pereira C, Pardo-Palacios FJ, Del Risco H, Ferrell M, Mellado M, Macchietto M, Verheggen K, et al: SQANTI: extensive characterization of long-read transcript sequences for quality control in full-length transcriptome identification and quantification. Genome Res 2018, 28:396–411.

48. Levin JZ, Berger MF, Adiconis X, Rogov P, Melnikov A, Fennell T, Nusbaum C, Garraway LA, Gnirke A: Targeted next-generation sequencing of a cancer transcriptome enhances detection of sequence variants and novel fusion transcripts. Genome Biol 2009, 10:R115.

49. Gnirke A, Melnikov A, Maguire J, Rogov P, LeProust EM, Brockman W, Fennell T, Giannoukos G, Fisher S, Russ C, et al: Solution hybrid selection with ultra-long oligonucleotides for massively parallel targeted sequencing. Nat Biotechnol 2009, 27:182–189.

50. Dixit A: Correcting Chimeric Crosstalk in Single Cell RNA-seq Experiments. bioRxiv 2021:093237.

51. Guanghao Q, Benjamin JS, Joshua MP, Hongkai J, Alexis B: Single-cell allele-specific expression analysis reveals dynamic and cell-type-specific regulatory effects. bioRxiv 2022:2022.2010.2006.511215.

52. Brooks ME, Kristensen K, van Benthem KJ, Magnusson A, Berg CW, Nielsen A, Skaug HJ, Mächler M, Bolker BM: glmmTMB Balances Speed and Flexibility Among Packages for Zero-inflated Generalized Linear Mixed Modeling. The R Journal 2017, 9.

53. Squair JW, Gautier M, Kathe C, Anderson MA, James ND, Hutson TH, Hudelle R, Qaiser T, Matson KJE, Barraud Q, et al: Confronting false discoveries in single-cell differential expression. Nat Commun 2021, 12:5692.

54. Crowell HL, Soneson C, Germain PL, Calini D, Collin L, Raposo C, Malhotra D, Robinson MD: muscat detects subpopulation-specific state transitions from multi-sample multi-condition single-cell transcriptomics data. Nat Commun 2020, 11:6077.

55. de Klein N, Tsai EA, Vochteloo M, Baird D, Huang Y, Chen CY, van Dam S, Oelen R, Deelen P, Bakker OB, et al: Brain expression quantitative trait locus and network analyses reveal downstream effects and putative drivers for brain-related diseases. Nat Genet 2023, 55:377–388.

56. Shabalin AA: Matrix eQTL: ultra fast eQTL analysis via large matrix operations. Bioinformatics 2012, 28:1353–1358.

57. Liu H, Prashant NM, Spurr LF, Bousounis P, Alomran N, Ibeawuchi H, Sein J, Slowinski P, Tsaneva-Atanasova K, Horvath A: scReQTL: an approach to correlate SNVs to gene expression from individual scRNA-seq datasets. BMC Genomics 2021, 22:40.

58. Wang AT, Shetty A, O’Connor E, Bell C, Pomerantz MM, Freedman ML, Gusev A: Allele-Specific QTL Fine Mapping with PLASMA. Am J Hum Genet 2020, 106:170–187.

59. Genovese G, Kahler AK, Handsaker RE, Lindberg J, Rose SA, Bakhoum SF, Chambert K, Mick E, Neale BM, Fromer M, et al: Clonal hematopoiesis and blood-cancer risk inferred from blood DNA sequence. N Engl J Med 2014, 371:2477–2487.

60. Jaiswal S, Fontanillas P, Flannick J, Manning A, Grauman PV, Mar BG, Lindsley RC, Mermel CH, Burtt N, Chavez A, et al: Age-related clonal hematopoiesis associated with adverse outcomes. N Engl J Med 2014, 371:2488–2498.

61. Robles-Espinoza CD, Mohammadi P, Bonilla X, Gutierrez-Arcelus M: Allele-specific expression: applications in cancer and technical considerations. Curr Opin Genet Dev 2021, 66:10–19.

62. Zou L, S., Zhao T, Cable D, M., Murray E, Aryee M, J., Chen F, Irizarry R, A.: Detection of allele-specific expression in spatial transcriptomics with spASE. bioRxiv 2021:2021.2012.2001.470861.

63. Beach TG, Adler CH, Sue LI, Serrano G, Shill HA, Walker DG, Lue L, Roher AE, Dugger BN, Maarouf C, et al: Arizona Study of Aging and Neurodegenerative Disorders and Brain and Body Donation Program. Neuropathology 2015, 35:354–389.

64. Habib N, Li Y, Heidenreich M, Swiech L, Avraham-Davidi I, Trombetta JJ, Hession C, Zhang F, Regev A: Div-Seq: Single-nucleus RNA-Seq reveals dynamics of rare adult newborn neurons. Science 2016, 353:925–928.

65. Volden R, Vollmers C: Single-cell isoform analysis in human immune cells. Genome Biol 2022, 23:47.

66. Wenger AM, Peluso P, Rowell WJ, Chang PC, Hall RJ, Concepcion GT, Ebler J, Fungtammasan A, Kolesnikov A, Olson ND, et al: Accurate circular consensus long-read sequencing improves variant detection and assembly of a human genome. Nat Biotechnol 2019, 37:1155–1162.

67. Li H, Handsaker B, Wysoker A, Fennell T, Ruan J, Homer N, Marth G, Abecasis G, Durbin R, Genome Project Data Processing S: The Sequence Alignment/Map format and SAMtools. Bioinformatics 2009, 25:2078–2079.

68. Zorita E, Cusco P, Filion GJ: Starcode: sequence clustering based on all-pairs search. Bioinformatics 2015, 31:1913–1919.

69. Li H: Minimap2: pairwise alignment for nucleotide sequences. Bioinformatics 2018, 34:3094–3100.

70. Danecek P, Bonfield JK, Liddle J, Marshall J, Ohan V, Pollard MO, Whitwham A, Keane T, McCarthy SA, Davies RM, Li H: Twelve years of SAMtools and BCFtools. Gigascience 2021, 10.

71. Pedersen BS, Quinlan AR: Mosdepth: quick coverage calculation for genomes and exomes. Bioinformatics 2018, 34:867–868.

72. De Coster W, D’Hert S, Schultz DT, Cruts M, Van Broeckhoven C: NanoPack: visualizing and processing long-read sequencing data. Bioinformatics 2018, 34:2666–2669.

73. Stuart T, Butler A, Hoffman P, Hafemeister C, Papalexi E, Mauck WM, 3rd, Hao Y, Stoeckius M, Smibert P, Satija R: Comprehensive Integration of Single-Cell Data. Cell 2019, 177:1888–1902 e1821.

74. Wolock SL, Lopez R, Klein AM: Scrublet: Computational Identification of Cell Doublets in Single-Cell Transcriptomic Data. Cell Syst 2019, 8:281–291 e289.

75. Hao Y, Hao S, Andersen-Nissen E, Mauck WM, 3rd, Zheng S, Butler A, Lee MJ, Wilk AJ, Darby C, Zager M, et al: Integrated analysis of multimodal single-cell data. Cell 2021, 184:3573–3587.

76. Nicol PB, Miller J, W.: Model-based dimensionality reduction for single-cell RNA-seq using generalized bilinear models. bioRxiv 2023:2023.2004.2021.537881.

77. Bais AS, Kostka D: scds: computational annotation of doublets in single-cell RNA sequencing data. Bioinformatics 2020, 36:1150–1158.

78. Zheng GX, Terry JM, Belgrader P, Ryvkin P, Bent ZW, Wilson R, Ziraldo SB, Wheeler TD, McDermott GP, Zhu J, et al: Massively parallel digital transcriptional profiling of single cells. Nat Commun 2017, 8:14049.

79. Cunningham F, Allen JE, Allen J, Alvarez-Jarreta J, Amode MR, Armean IM, Austine-Orimoloye O, Azov AG, Barnes I, Bennett R, et al: Ensembl 2022. Nucleic Acids Res 2022, 50:D988–D995.

80. R: A Language and Environment for Statistical Computing [https://www.R-project.org/]

81. Townes FW, Hicks SC, Aryee MJ, Irizarry RA: Feature selection and dimension reduction for single-cell RNA-Seq based on a multinomial model. Genome Biol 2019, 20:295.

82. Korsunsky I, Millard N, Fan J, Slowikowski K, Zhang F, Wei K, Baglaenko Y, Brenner M, Loh PR, Raychaudhuri S: Fast, sensitive and accurate integration of single-cell data with Harmony. Nat Methods 2019.

83. Traag VA, Waltman L, van Eck NJ: From Louvain to Leiden: guaranteeing well-connected communities. Sci Rep 2019, 9:5233.

84. McInnes L, Healy J, Melville J: UMAP: Uniform Manifold Approximation and Projection for Dimension Reduction. arXiv 2020:1802.03426.

85. Finak G, McDavid A, Yajima M, Deng J, Gersuk V, Shalek AK, Slichter CK, Miller HW, McElrath MJ, Prlic M, et al: MAST: a flexible statistical framework for assessing transcriptional changes and characterizing heterogeneity in single-cell RNA sequencing data. Genome Biol 2015, 16:278.

86. Quinlan AR, Hall IM: BEDTools: a flexible suite of utilities for comparing genomic features. Bioinformatics 2010, 26:841–842.

87. Di Tommaso P, Chatzou M, Floden EW, Barja PP, Palumbo E, Notredame C: Nextflow enables reproducible computational workflows. Nat Biotechnol 2017, 35:316–319.

88. Liao Y, Smyth GK, Shi W: featureCounts: an efficient general purpose program for assigning sequence reads to genomic features. Bioinformatics 2014, 30:923–930.

89. Smith T, Heger A, Sudbery I: UMI-tools: modeling sequencing errors in Unique Molecular Identifiers to improve quantification accuracy. Genome Res 2017, 27:491–499.

90. McKinney W: Data structures for statistical computing in python. In Proceedings of the 9th Python in Science Conference. Volume 445. Edited by van der Walt Se, Millman J; 2010: 56–61

91. Huang X, Huang Y: Cellsnp-lite: an efficient tool for genotyping single cells. Bioinformatics 2021, 37:4569–4571.

92. Cotto KC, Feng YY, Ramu A, Richters M, Freshour SL, Skidmore ZL, Xia H, McMichael JF, Kunisaki J, Campbell KM, et al: Integrated analysis of genomic and transcriptomic data for the discovery of splice-associated variants in cancer. Nat Commun 2023, 14:1589.

93. Johnson WE, Li C, Rabinovic A: Adjusting batch effects in microarray expression data using empirical Bayes methods. Biostatistics 2007, 8:118–127.

94. Leek JT, Johnson WE, Parker HS, Jaffe AE, Storey JD: The sva package for removing batch effects and other unwanted variation in high-throughput experiments. Bioinformatics 2012, 28:882–883.

95. Zheng X, Levine D, Shen J, Gogarten SM, Laurie C, Weir BS: A high-performance computing toolset for relatedness and principal component analysis of SNP data. Bioinformatics 2012, 28:3326–3328.

96. Obenchain V, Lawrence M, Carey V, Gogarten S, Shannon P, Morgan M: VariantAnnotation: a Bioconductor package for exploration and annotation of genetic variants. Bioinformatics 2014, 30:2076–2078.

97. Carithers LJ, Ardlie K, Barcus M, Branton PA, Britton A, Buia SA, Compton CC, DeLuca DS, Peter-Demchok J, Gelfand ET, et al: A Novel Approach to High-Quality Postmortem Tissue Procurement: The GTEx Project. Biopreserv Biobank 2015, 13:311–319.

